# Discovery of a redox-thiol switch regulating cellular energy metabolism

**DOI:** 10.1101/520411

**Authors:** Xing-Huang Gao, Ling Li, Marc Parisien, Matt Mcleod, Jing Wu, Ilya Bederman, Zhaofeng Gao, Dawid Krokowski, Steven M Chirieleison, Luda Diatchenko, Derek Abbott, Vivien Yee, Charles L. Hoppel, Richard G Kibbey, Todd Holyoak, Belinda Willard, Peter Arvan, Maria Hatzoglou

## Abstract

Previously, we reported that increased synthesis of the gas hydrogen sulfide (H_2_S) during the Integrated Stress Response (ISR) induced proteome-wide cysteine-sulfhydration with the predominant modified pathway being enzymes of cellular energy metabolism (Gao, et al. 2015). Using pancreatic beta cells and quantitative proteomics in this study, we identified a Redox Thiol Switch from S-glutathionylation to S-sulfhydration and we named it, RTS^GS^. About half of the identified proteins are involved in energy metabolism, and one novel target was the mitochondrial phosphoenolpyruvate carboxykinase 2 (PCK2) whose catalytic Cys^306^was targeted by both modifications. The enzymatic activity of PCK2 was inhibited by S-glutathionylation, and this inhibition was largely reversed by S-sulfhydration. S-sulfhydration also reversed the S-glutathionylation-mediated inhibition of glucose flux, indicating a broad metabolic significance. We propose that a Redox Thiol Switch from S-glutathionylation to S-sulfhydration is a key mechanism to fine tune cellular energy metabolism in response to different levels of oxidative stress.

## Introduction

Hydrogen sulfide (H_2_S) is a gas molecule that can be produced endogenously in many organisms from bacteria to mammals. H_2_S production at physiological concentrations is cytoprotective, as it reduces blood pressure (Yang et al., 2005), prevents neurodegeneration and extends lifespan (Hine et al., 2015; Kabil et al., 2011; Miller and Roth, 2007; Paul et al., 2014). H_2_S reacts with and modifies protein cysteine thiol groups to form persulfide bonds, known as protein S-sulfhydration. Because H_2_S is unstable, and its synthesis is regulated (Sbodio et al., 2018), protein S-sulfhydration serves as a mechanism to regulate protein functions with an impact in cellular metabolism (Mustafa et al., 2009, Gao et al., 2015).

During oxidative stress Reactive Oxygen Species (ROS) often catalyze the non-enzymatic protein S-sulfhydraton (Mishanina et al., 2015; Paul and Snyder, 2015). In normal cells, in the absence of oxidative stress, protein S-sulfhydration can be the result of either local subcellular ROS fluctuations, or the enzymatic local H_2_S synthesis (Paul and Snyder, 2015). The latter likely involves the shunting of the SH group of the H_2_S to protein thiol groups, by the H_2_S-producing enzymes, as previously suggested for protein S-nitrosylation (Paul and Snyder, 2015). Among all amino acid residues in proteins, cysteine is the most susceptible to oxidative modifications in which a sulfur group is covalently bound to a ROS-derived electrophilic molecule (Reddie and Carroll, 2008). Depending on the types and levels of ROS, a cysteine residue in a protein can be subjected to multiple types (Paul and Snyder, 2015) of redox-based posttranslational modifications (redox PTM). In addition to the already mentioned S-sulfhydration, other regulatory modifications are S-nitrosylation and S-glutathionylation (Gao et al., 2015, (Martinez-Ruiz and Lamas, 2007). S-nitrosylation is the covalent attachment of a nitroso group (-NO) to cysteine thiol of a protein to form an S-nitrosothiol (SNO). S-glutathionylation occurs when a protein cysteine residue forms an intermolecular disulfide bond with a glutathione to form protein-SSG.

There is a growing body of evidence that the switch from one redox PTM to another in proteins, plays an important role in the regulation of various biological functions (Paul and Snyder, 2015). More importantly, various types of redox PTMs at the same cysteine residue in a protein yield diverse effects on the functions of the given protein. For instance, S-glutathionylation of a single redox sensitive cysteine in GAPDH inhibits its enzymatic activity, while S-sulfhydration can reverse this inhibitory effect and restore its activity (Gao et al., 2015). Similarly, S-nitrosylation of the p65 subunit of NF-*kB* inhibits its DNA binding activity, but S-sulfhydration increases the association of NF-kB with promoters of cytoprotective target genes in response to inflammation (Sen et al., 2012). Nonetheless, the ratio of different modifications on the same protein in a given cellular context determines the biological outcome. We focused on S-glutathionylation which is a pathophysiological redox protein regulation mechanism through the local subcellular fluctuations of GSH and ROS levels (Mieyal et al., 2008). The presence and extent of the conversion of S-glutathionylation to other oxidative PTMs in the proteome, is largely unknown. Furthermore, in the case of the conversion of S-glutathionylation to S-sulfhydration, it is not clear if there is a switch of one modification to another on the same cysteine residue. The physiological consequences for such a protein Redox Thiol Switch (RTS) are also not well understood.

It has been recognized as a great challenge to detect and quantify protein S-sulfhydration ***in vivo,*** due to the high reactivity and instability of the cysteine persulfide bond in sulfhydrated proteins (Mishanina et al., 2015; Paul and Snyder, 2015). Earlier, we developed a proteomics approach (Gao et al., 2015), the Biotin Thiol Assay (BTA) to identify sylfhydrated proteins. We used this approach to show that the increase of H_2_S synthesis during endoplasmic reticulum (ER) stress in pancreatic beta cells promotes selective S-sulfhydration of redox sensitive cysteine residues in proteins of specific metabolic pathways (Gao et al., 2015). The previous studies implicated H_2_S as a molecular switch in the regulation of energy metabolism.

In this current study, we show the existence of an RTS, which involves H_2_S-mediated reversal of S-glutathionylation to S-sulfhydration (RTS^GS^) of specific cysteine residues in a subset of glutathionylated proteins. The iodoacetyl isobaric tandem mass tag system (iodoTMT) was combined with the previously reported (Gao et al., 2015) BTA assay (TMT-BTA), thus enabling the quantitative identification of sulfhydrated or RTS^GS^ target proteins. We identified distinct patterns of the natural protein sulfhydromes in human pancreatic beta cells and hepatocytes, with hepatocytes exhibiting higher levels of sulfhydrated proteins compared to pancreatic beta cells. However, human pancreatic beta cells exhibited a unique landscape in sulfhydrated proteins, with the most enriched pathways being in intermediary metabolism. Furthermore, we identified insulin, the beta cell specific protein and master regulator of cellular metabolism, as a S-sulfhydration target. A novel RTS^GS^ target in beta cells was the metabolic mitochondrial enzyme phosphoenolpyruvate carboxykinase 2 (PCK2) and known regulator (Stark et al., 2009) of glucose-stimulated insulin secretion (GSIS). The catalytic residue Cys^306^ was either sulfhydrated or glutathionylated. *In vitro* biochemical characterization showed that S-glutathionylation of the human recombinant PCK2 protein inhibited its enzymatic activity, whereas S-sulfhydration reversed the inhibitory effect of S-glutathionylation on PCK2 activity. Finally, in order to understand the physiological significance of this RTS^GS^ in cellular energy metabolism, we induced protein S-glutathionylation in pancreatic beta cells treated with diamide and measured metabolic flux of glucose. It is well known that protein S-glutathionylation can inhibit the activities of metabolic enzymes (Mieyal et al., 2008), and cause a decrease in the metabolic flux of glucose (Hiranruengchok and Harris, 1995). In agreement with the RTS^GS^ mechanism contributing to regulation of energy metabolism, H_2_S, rescued the inhibited glucose flux in cells treated with diamide. We conclude that RTS^GS^, the redox thiol switch from S-glutathionylation to S-sulfhydration is a mechanism to fine tune cellular energy metabolism in response to oxidative stress.

## Results

We have used the previously reported by us, BTA assay (Gao et al., 2015), to develop a quantitative proteomics approach for profiling of the changes in the protein sulfhydrome in different cellular contexts. The utilization of tandem mass tags (TMT) is a well-recognized proteomics approach for the relative quantification of proteins and/or peptides (Thompson et al., 2003). We therefore have included the process of iodoTMT multiplex labeling of the peptides in the final step of the BTA assay (Fig 1A). The TMT-labeled sulfhydrated peptides can be combined for LC-MS identification and quantification.

**Figure 1.**
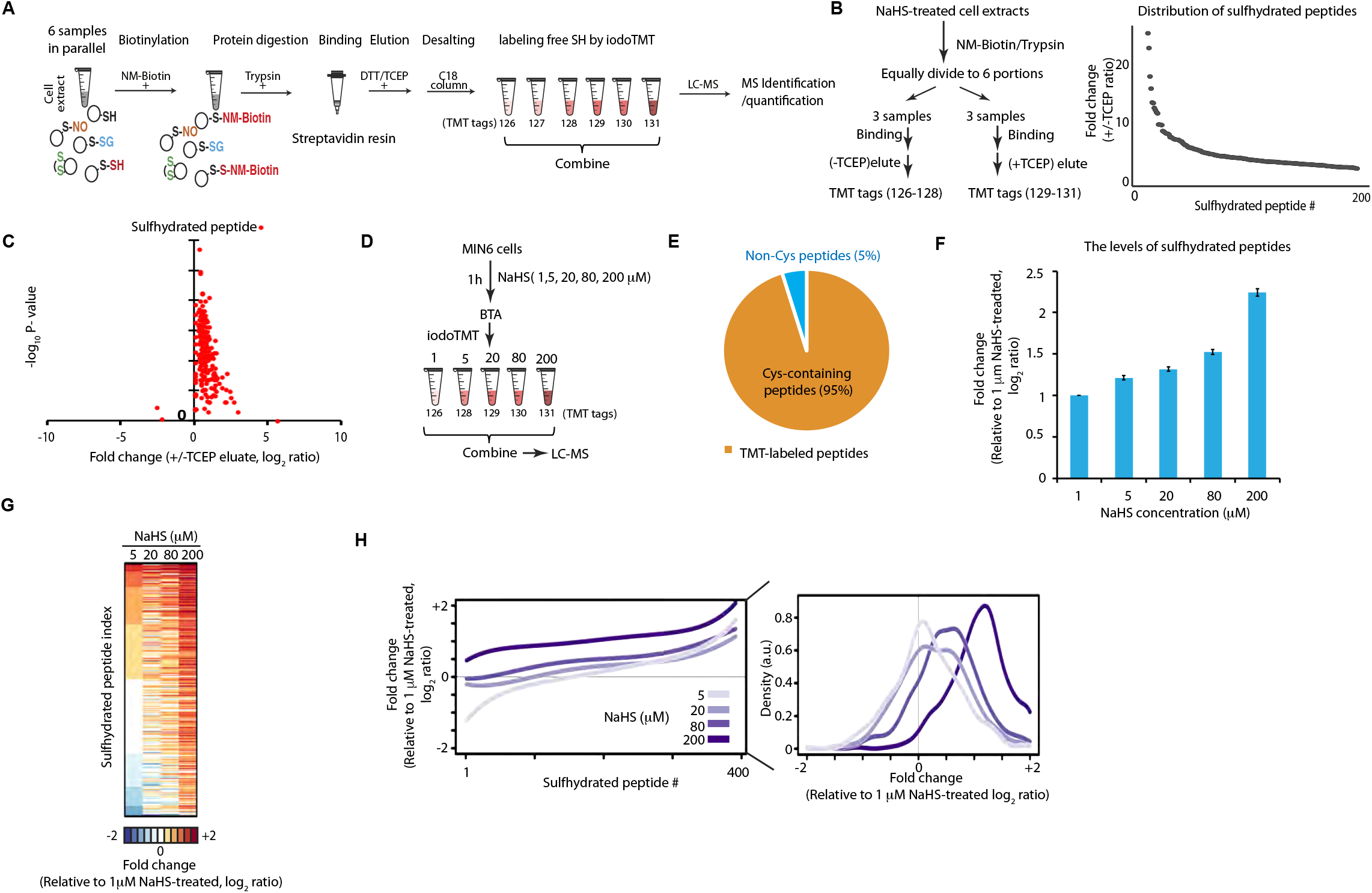
Quantitative proteomics approach to profile the protein sulfhydrome *in vivo.* **(A)** Graphic representation of the Biotin Thiol Assay (BTA) (Gao et al., 2015) combined with alkylation of free SH groups with the tandem mass tags sixplex system (iodoTMT). Sulfhydrated cysteine residues in proteins from 6 samples were biotinylated with maleimide-PEG2-biotin (NM-Biotin). Subsequently, samples were processed with trypsin, passed through an avidin column and eluted with DTT or TCEP, which reduced the persulfide bonds and produced eluates containing the peptides with free SH groups. After desalting, free SH groups were alkylated with iodoTMT tags and the TMT-labeled samples were combined for LC-MS analysis. **(B)** Schematic of workflow to determine the specificity of the TMT-BTA assay *(left panel).* Protein extracts from mouse pancreatic beta cells (MIN6) pretreated with an H_2_S donor (NaHS, 0.2 mM) were lysed, then subjected to the TMT-BTA assay. After trypsin digestion of the biotinylated proteome, the peptide sample was divided into six equal fractions. Each fraction was incubated with the same amount of avidin beads, then eluted without (-) or with (+) TCEP followed by the TMT labeling. The TMT-labeled samples were combined and subjected to LC-MS. Curve plot showing the distribution of relative changes of ratios (+/-TCEP) of the sulfhydrated peptides against the number (#) of identified peptides *(right panel).* Triplicates of each condition were used **(C)** Volcano plot showing the *P* value distribution against the relative changes of ratios (+/-TCEP) of the sulfhydrated peptides identified from the experimental data in (B). **(D)** Schematic workflow to evaluate the TMT-BTA assay for the quantification of sulfhydrated proteins *in vivo.* MIN6 cells were treated with the H_2_S donor, NaHS, at the indicated concentrations for 1h, then subjected to the TMT-BTA assay. LC-MS analysis was used to identify and quantify sulfhydrated peptides. **(E)** Pie chart illustrating that 95% of the identified peptides in (D), were containing cysteine residues. **(F)** Bar plot illustrating the dose-dependent increases of the average values of relative ratios (log_2_) of sulfhydrated peptides identified in (D). **(G)** Heat map showing the relative sulfhydrated peptide enrichment ratios in MIN6 cells treated with the H_2_S donor at the indicated concentrations, as shown in (D). **(H)** *(Left panel)* Curve plot showing the distribution of relative sulfhydrated peptide enrichment ratios against the number of identified peptides from the experimental data in (D), *(Right panel)* a histogram chart for the left panel.

We first optimized the buffer system and the concentration of reducing agents (DTT or TCEP) in the elution buffer of the BTA assay, in order to maximize TMT-labeled peptides in the eluate. We treated mouse pancreatic beta cells (MIN6) with a commonly used H_2_S donor (NaHS, sodium hydrosulfide), which promotes protein S-sulfhydration both *in vitro* and *in vivo* (Paul and Snyder, 2015). The NaHS-treated MIN6 cells were lysed and subjected to the BTA assay. The sulfhydrated peptides were eluted in different buffer systems (A-D, Supplemental Fig 1) followed by the iodoTMT labeling and LC-MS analysis (Supplemental Fig 1). The elution buffer containing 50 mM TEAB (triethylammonium bicarbonate) and 10 mM TCEP (tris-carboxyethyl phosphine hydrochloride) provided the highest labeling efficiency of sulfhydrated peptides (Supplemental Fig 1). As a first step, in order to identify the range of protein targets in the entire cellular proteome that are potentially subjected to RTS^gs^ in living cells, we treated MIN6 cell lysates with NaHS and then applied the optimized TMT-BTA assay (Fig 1B, left panel). Elution was performed either with a buffer containing TCEP, or NaCl (-TCEP) as a negative control. Eluates were labeled with iodoTMT reagents, followed by LC-MS analysis (Fig 1B, *left panel*). About 200 sulfhydrated peptides were identified and quantified (Fig 1B, *right panel*). Compared to the NaCl control, TCEP-eluted samples exhibited significantly higher ratios of sulfhydrated peptides (Fig 1C). These data verify that the TMT-BTA assay is a highly selective method to identify the protein sulfhydromes *in vitro.*

We next determined the changes in the cellular sulfhydrome *in vivo* under exposure to increasing H_2_S levels. The intracellular levels of H_2_S are on the order of nanomolar to micromolar, depending on cell types and cellular contexts (Filipovic et al., 2018). Under the highly reducing intracellular environments, protein S-sulfhydration is unlikely the predominant modification over the vast majority of protein free thiol groups in the proteome. Therefore, to determine if the TMT-BTA assay can quantify stoichiometric changes in the protein sulfhydrome in cells, we incubated MIN6 cells with increasing concentrations of the H_2_S donor (NaHS, 1-200 μM) followed by the TMT-BTA of MIN6 cell extracts (Fig 1D). The vast majority (95%) of the identified peptides by the TMT-BTA assay contained cysteine residues, confirming that this method is highly selective in the identification of cysteine persulfides (Fig 1E). As expected, the increase in the levels of protein S-sulfhydration was dependent on the NaHS concentration (Fig 1F-H). Moreover, we found that all MS-identified sulfhydrated peptides were labeled by the TMT tags (data not shown). Taken together, these data show that the TMT-BTA assay is a tool to quantitatively study changes of the S-sulfhydrated proteome in live cells and cell extracts under different redox manipulations.

Next we determined the ability of the TMT-BTA assay to distinguish the levels of sulfhydromes between different cell types. It has been reported that protein S-sulfhydration is a prevalent redox PTM in certain tissues; for example, about 10–25% of mouse hepatic proteins are estimated to be sulfhydrated (Mustafa et al., 2009). The high abundance of protein S-sulfhydration in liver cells made this cell type ideal for our purpose of comparing sulfhydromes between cell types. To this aim, we used the TMT-BTA assay to analyze the protein sulfhydrome in immortalized human hepatocytes, IHH, (Ulatowski et al., 2012) and in human beta cells, EndoC-BH3 (Fig 2A). 447 sulfhydrated peptides from 307 proteins were identified and quantified (Fig 2B). Among all identified proteins, 33 of them were sulfhydrated specifically in EndoC-BH3 cells with ratios greater than 1.5-fold (Fig 2C-D). As expected, human hepatocytes exhibited significantly higher levels of sulfhydrated peptides as compared to EndoC-BH3 cells (Fig 2B-D). Bioinformatics clustering with a pathway annotation program (DAVID) of the identified shared sulfhydrome between hepatocytes and EndoC-BH3 cells, revealed an enrichment of those proteins involved in ribosome and metabolic pathways including glycolysis/gluconeogenesis (Fig 2E). Furthermore, proteins involved in mRNA translation and metabolic pathways are highly susceptible to S-sulfhydration, in agreement with our previous studies showing that the stress-induced inhibition of glycolytic flux was reversed by increasing H_2_S biosynthesis in the presence of stress (Gao et al., 2015). The higher levels of protein S-sulfhydration in hepatocytes as compared to the EndoC-BH3 cells, was further supported by the higher expression levels of the H_2_S-generating enzymes in these cells (Fig 2F). De novo synthesis of H_2_S is mediated by three enzymes: CTH (cystathionine γ lyase), CBS (cystathionine β synthase) and 3-MST (3-mercaptopyruvate sulfurtransferase) (Sbodio et al., 2018). Therefore, the levels of sulfhydrated proteins are in good correlation with the levels of the H_2_S-generating enzymes in these cells (Fig 2).

**Figure 2.**
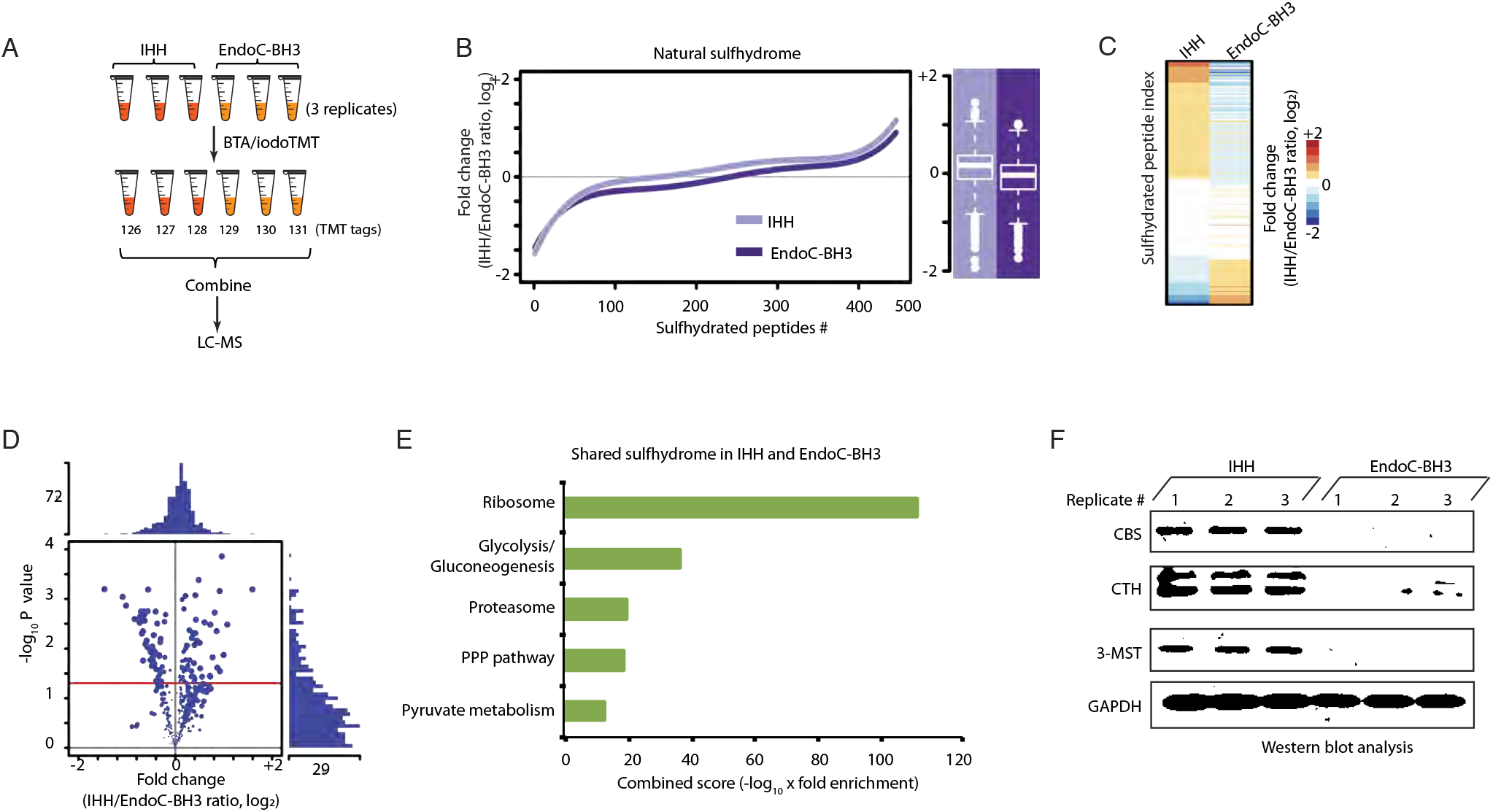
Quantitative profiling of the natural protein sulfhydrome in human hepatocytes and pancreatic beta cells by the TMT-BTA assay. **(A)** Schematic of workflow for profiling the sulfhydrome in the two cell lines. Protein extracts from human hepatocytes (IHH) and pancreatic beta cells (EndoC-BH3) were subjected to the TMT-BTA assay, and the TMT-labeled peptides were mixed and analyzed by LC-MS. Triplicates of each cell line were used. **(B)** *(Left panel)* Curve plot showing the distribution of relative changes (log_2_ IHH/EndoC-BH3 ratios) of the sulfhydrated peptides in IHH (light purple) and EndoC-BH3 cells (deep purple). *(Right panel)* Boxplot showing comparisons of the average levels of relative ratios (log_2_) of sulfhydrated peptides between the two cell lines. Triplicates of each cell line were used. **(C)** Heat map of the relative ratios of the sulfhydrated peptides identified in experimental data in (B), illustrating the differences in the patterns of the natural protein sulfhydrome between IHH and EndoC-BH3 cells. **(D)** Volcano plot showing the *P* value distribution against the fold change of ratios (log_2_ IHH/EndoC-BH3 ratios) of the sulfhydrated peptides identified from the experimental data in (B). **(E)** Gene ontology biological pathways for all sulfhydrated proteins. PPP: pentose phosphate pathway. **(F)** Evaluation of levels of H_2_S-generating proteins (CTH, CBS and 3-MST) in IHH and EndoC-BH3 by Western blot analysis.

We hypothesized that the shared hepatic and beta cell sulfhydrome overlaid on their proteomes should show differential pathway enrichments. We first performed a label-free MS quantification experiment to determine the absolute protein abundance in the cellular proteomes (Fig 3A). With this analysis, almost 4500 proteins were identified and their abundance was quantified and compared between the two cell lines (Fig 3B). About 2-3% of these proteins were solely identified either in the EndoC-BH3 cells, or in the hepatocytes; 10-13% of the proteins exhibited 2-fold differences between the two cell types, including CBS and CTH, which were in agreement with the Western blot analyses shown in Fig 2F. To further validate the label-free MS data, we used Western blot assays to test the expression of two proteins, MANF (mesencephalic astrocyte-derived neurotrophic factor) and PCK2 (mitochondrial phosphoenolpyruvate carboxykinase), which exhibited differential protein abundance in the two cell lines. In agreement with the MS data, EndoC-BH3 cells had higher expression of the MANF protein (Supplemental Fig 3). In contrast, no significant change was found in the metabolic enzyme PCK2 levels between hepatocytes and EndoC-BH3 cells, in agreement with the MS data (Fig 3B). Pathway analysis of the proteins with relative abundance greater than 2-fold between the two cell types (Fig 3B), revealed that the branched-chain amino acid degradation and ribosome pathways were enriched in EndoC-BH3 cells and hepatocytes, respectively (Fig 3D). Next, we compared the relative changes in sulfhydromes between the two cell types, over their proteome abundance. To this end, we overlaid the protein abundance map with the natural sulfhydrome datasets (Fig 3E). This analysis showed that 63 sulfydrated proteins had a relative high abundance (log2 ratios greater than 1) in hepatocytes and 32 sulfhydrated proteins had relative high abundance (log2 ratios less than −1) in EndoC-BH3 cells. Among the 62 proteins, the enriched pathways were the ribosome and gluconeogenesis/glycolysis, whereas no pathway enrichment was noted in the 32 proteins from the EnoC-BH3 cells. Furthermore, the remaining sulfhydrated proteins identified in both cell types showed the proteasome as an enriched pathway. Together, these data suggest that cellular sulfhydromes exhibit differential pathway enrichments as compared to their proteomes. A few metabolic proteins, such as lactase dehydrogenase (LDHA and LDHB), fructose-bisphosphate aldolase (ALDOA) and enoase 1 (ENO1) (Fig 3F), were in relatively lower abundance, but were more sulfhydrated in the EndoC-BH3 cells than the hepatocytes. Overall, the evaluation of the relative abundance of sulfhydromes described here, can also be a very useful tool for identifying changes in the redox-mediated cellular responses in a single cell type under different intracellular and extracellular cues.

**Figure 3.**
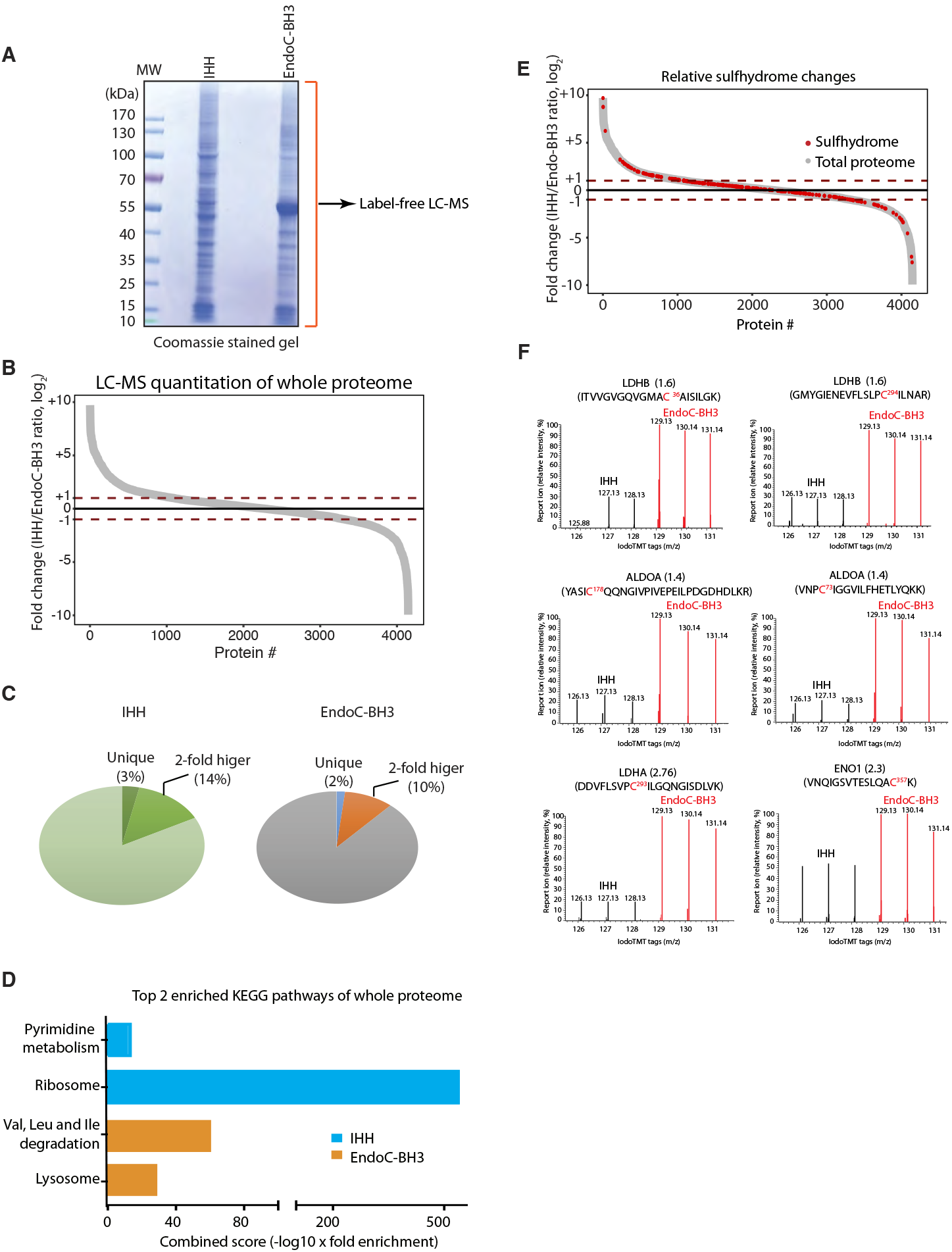
Protein S-sulfhydration does not correlate with relative protein abundance between IHH and EndoC-BH3 cells. **(A)** Identification and quantification of the full proteome from human hepatocytes (IHH) and EndoC-BH3 cells by LC-MS analysis. Cell extracts from both cell lines were resolved by reducing SDS–PAGE and stained with Coomassie blue. Gel lane was cut into fractions for in-gel digestions with trypsin. The peptides from each fraction were submitted for LC-MS identification and quantification. A total of 4664 proteins from hepatocytes and 4050 proteins from EndoC-BH3 cells were identified and quantified by LC-MS. **(B)** Comparison of protein abundance between human hepatocytes and EndoC-BH3 cells in a curve plot showing the cell-type specific differences. Red dotted lines marks the sulfhydration ratio (IHH/EndoC-BH3, log_2_) greater or less than 1. **(C)** Pie charts of the data described in A and B. **(D)** Gene ontology biological pathways for the relative most abundant proteins (> 1 in Fig 3B) identified from IHH (blue bars) and EndoC-BH3 cells (orange bars). **(E)** Comparison of the natural protein sulfhydrome in Fig 2B with the full proteome reveals that a large fraction of sulfhydrated proteins have a moderate abundance in the cells. **(F)** The TMT-BTA profiling of proteins containing cysteine residues with different reactivity to H_2_S from IHH and EndoC-BH3 cells. LC-MS-MS(2) profiles for cysteine-containing peptides from LDHB, ALDOA, LDHA and ENO1 proteins are shown.

Next, we used the TMT-BTA assay to determine the regulation of the sulfhydrome in a single cell line under genetic manipulation of de novo H_2_S synthesis. We chose INS1 cells because the levels of the H_2_S-producing enzyme CTH, was undetectable in the absence of ER stress (Fig 4E) and thus made this cell line a good candidate to determine the effects of CTH expression on the sulfhydrome. We have previously shown that during chronic ER stress in MIN6 cells, increased protein S-sulfhydration was in good correlation with increased expression of the transcription factor ATF4 and its transcription target, the CTH gene (Sbodio et al., 2018). In order to get more insights on the regulation of the sulfhydromes by the ATF4/CTH axis-mediated de novo H_2_S synthesis (Fig 4A), we overexpressed the transcription factor ATF4 or GFP (green fluorescent protein) as a negative control in INS1 cells and applied the TMT-BTA assay to cell lysates. The TMT-BTA assay identified and quantified over 2000 sulfhydrated peptides (Fig 4B). Cysteine S-sulfhydration levels were significantly increased by the forced ATF4 expression but not by GFP expression (Fig 4B-C), in agreement to our previous study (Gao et al., 2015). However, ATF4 can induce expression of many genes that can potentially increase the sulfhydrome (Gao et al., 2015; Mieyal et al., 2008). To identify the subgroup of CTH-mediated sulfhydrated proteins (Fig 4A), we looked for proteins with significant decreases in S-sulfhydration in response to depletion of CTH in ATF4-expressing INS1 cells. To this end, we employed the CRISPR-Cas9 technique to knock out the CTH-encoding gene in INS1 cells. We designed two single guide RNA sequences to target the translation start codon of the rat CTH gene (Fig 4D, Supplemental Fig 4-1A), and then screened for individual INS1 clones to identify the deficiency of CTH expression induced by ATF4 (Fig 4E, Supplemental Fig 4-1(B, C, D) and Supplemental Fig 4-2). INS1 cells deficient for CTH, were infected with adenovirus that expressed ATF4, followed by the TMT-BTA assay. As expected, CTH depletion resulted in significant decreases of the protein sulfhydrome in INS1 cells expressing ATF4 (Fig 4F-I), confirming that regulation of protein S-sulfhydration by ATF4, is mediated in part via the induction of CTH. Among about 2000 sulfhydrated peptides, 157 peptides (corresponding to 130 proteins) exhibited more than 1.5-fold decrease in S-sulfhydration with the loss of CTH induction. Notably, insulin exhibited a significant CTH-dependent decrease in S-sulfhydration, suggesting that insulin may be a specific substrate of CTH in INS1 cells expressing ATF4. Western blot analysis using antibodies for proinsulin revealed that the protein expression levels were not affected by CRISPR-mediated CTH knockout in INS1 cells expressing ATF4 (Fig 4J), further supporting that the decreased insulin S-sulfhydration is not due to changes in intracellular insulin content.

**Figure 4.**
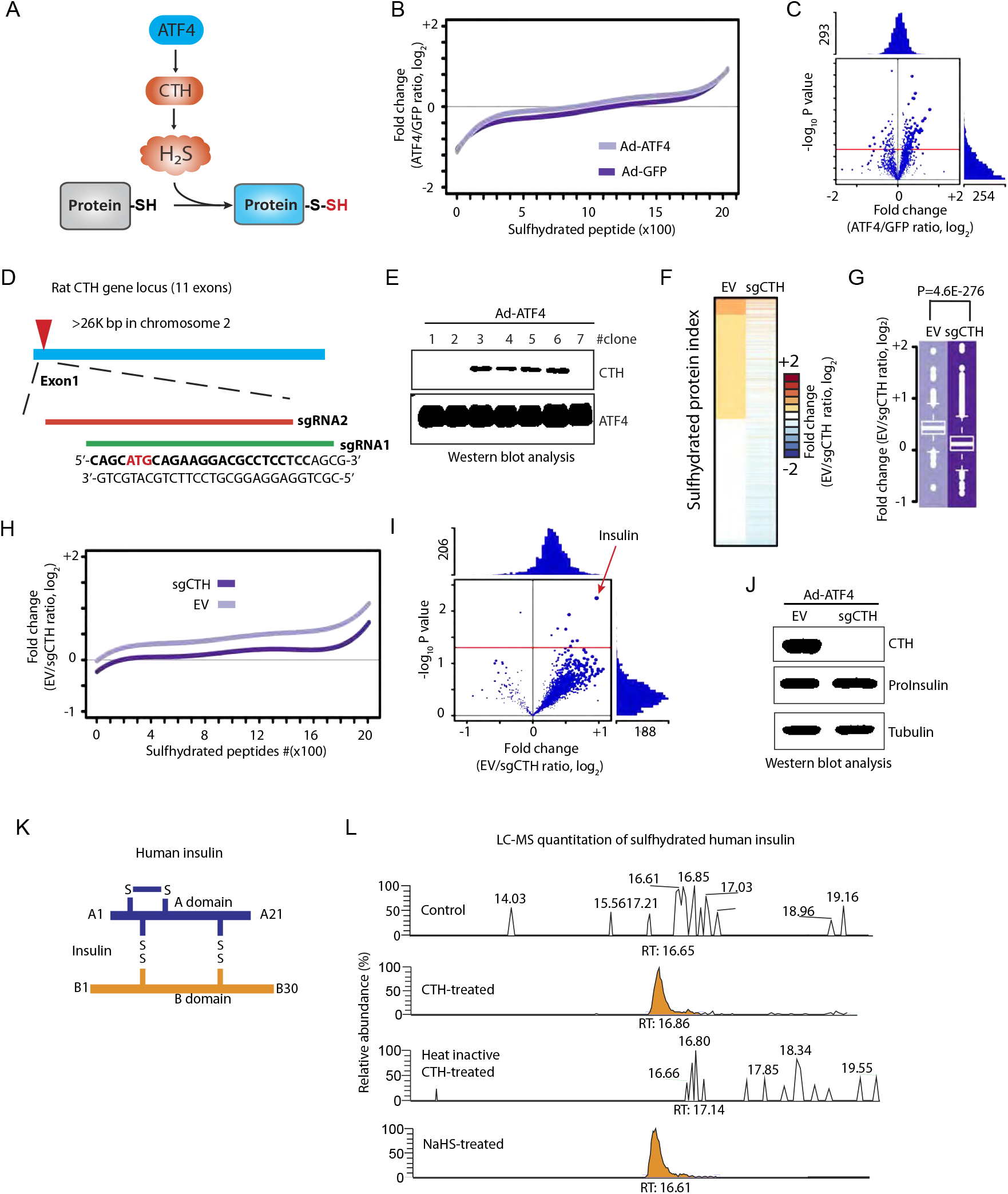
The TMT-BTA assay reveals insulin as a CTH-mediated sulfhydrated protein, in INS1, rat pancreatic beta cells. **(A)** Schematic representation of the ATF4-induced CTH gene expression leading to increased intracellular H_2_S synthesis and S-sulfhydration of proteins. **(B)** Relative distribution of peptides containing sulfhydrated cysteine residues as determined by the TMT-BTA assay of cell extracts isolated from INS1 cells overexpressing ATF4 or GFP for 36 h. The relative changes (log_2_ value) of TMT ratios are plotted against the number of identified peptides. (C) Volcano plot showing the relative change of the sulfhydration ratios (ATF4/GFP, log_2_) of modified peptides plotted against the –log_10_ *P* value (n=3). Red line indicates the *P* value ≤ 0.05. **(D)** Schematic representation of CRISPR-cas9 mediated CTH knockout (KO) in INS1 cells. Two sgRNA sequences were designed for targeting exon 1 of the rat CTH gene. sgRNA targeting sequences are in bold. The translation initiation codon is shown in red. **(E)** Western blot analysis of the effect of ATF4-overexpression in individual CRISPR-mediated CTH knockout INS1 cells. Clones 1, 2 and 7 did not express CTH protein. **(F)** Heat map showing the relative enrichment values (log_2_ fold change) of sulfhydrated peptides identified in CTH KO (sgCTH) and control INS1 cells (empty vector, EV). CTH KO and EV INS1 cells were transfected to express ATF4, then subjected to the TMT-BTA assay, and followed by LC-MS identification and quantification. INS1 cells expressing cas9 protein without sgRNA were used as control. Triplicates of each condition were used. **(G)** Boxplot showing comparisons of the average values of the log_2_ ratio of sulfhydrated peptides from CTH KO and EV INS1 cells expressing ATF4. **(H)** Relative distributions of peptides containing sulfhydrated cysteine residues to their relative log_2_ values determined by the TMT-BTA assay. The relative changes (log_2_ value) of TMT ratios are plotted against the number of identified peptides from the experimental data in **(F). (I)** Volcano plot showing the log_2_ fold change (sgCTH versus EV) of sulfhydrated proteins against the –log_10_ *P* value. Insulin is a major sulfhydrated protein regulated by the CTH protein levels. **(J)** Western blot analysis of intracellular proinsulin levels showing no significant change of the levels of the protein in sgCTH INS1 cells as compared to the control cells. **(K)** Schematic representation of insulin inter/intra-molecular disulfides. **(L)** LC-MS analysis of CTH-mediated insulin sulfhydration or NaHS-treated insulin *in vitro.* Human recombinant insulin was incubated with active or heat-inactivated recombinant CTH in the presence of PLP and L-cysteine for 1h. The reaction mixture was subjected to LC-MS for the identification of sulfhydrated insulin. PLP: pyridoxal phosphate-a cofactor for the CTH enzyme. RT: retention time.

There are three disulfide bonds that are essential for the mature insulin to maintain the correct structure (Fig 4K). To verify whether the enzymatic activity of CTH can regulate insulin S-sulfhydration *in vitro,* we incubated recombinant human insulin with either active, or heat-inactivated recombinant CTH protein, followed by LC-MS analysis. Consistent with the *in vivo* data from INS1 cells, human insulin was sulfhydrated by H_2_S generated from the human recombinant CTH, but this modification was abolished by the inactivation of the CTH enzyme (Fig 4L). Treatment of recombinant insulin with NaHS gave similar results (Fig 4L). The combined data suggest that insulin S-sulfhydration is the result of an enzymatic reaction that requires the presence of active CTH to catalyze protein cysteine persulfidation.

We next tested the hypothesis that S-sulfhydration may be used as a regulatory mechanism of the enzymatic activities of metabolic proteins under oxidative stress conditions. It is known that redox sensitive cysteine glutathionylation inhibits the activities of metabolic enzymes (Mieyal et al., 2008). We have also previously shown that S-glutathionylation of GAPDH at the catalytic cysteine residue inhibits its enzymatic activity, and it can be restored by H_2_S treatment (Gao et al., 2015), suggesting that the S-glutathionylation to S-sulfhydration RTS switch (Fig 5A) could be a regulatory mechanism of energy metabolism.

**Figure 5.**
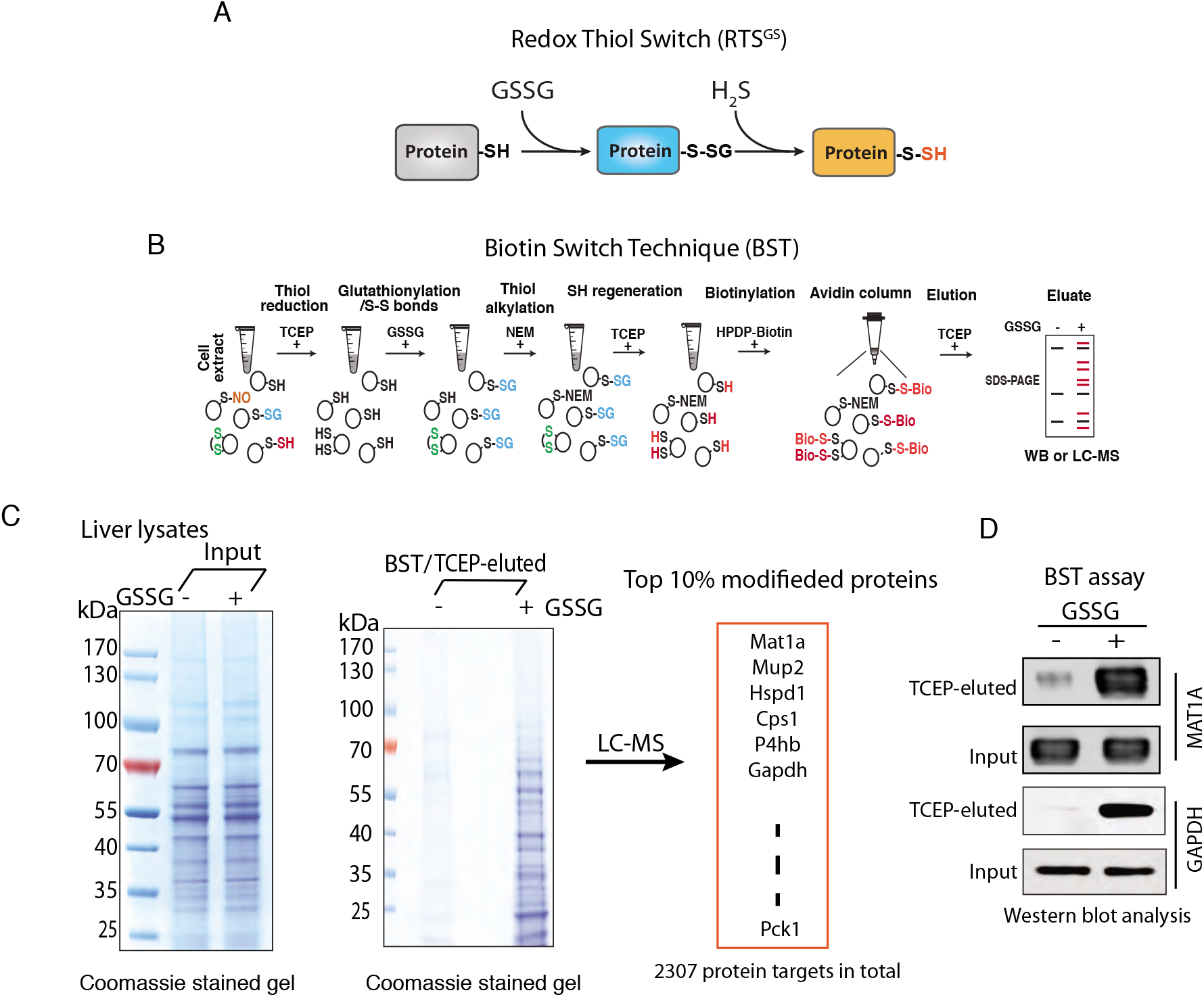
A proteomics approach to detect the RTS^GS^ switch *in vitro.* **(A)** Schematic representation of the Redox Thiol Switch (RTS^GS^) from S-glutathionylation to S-sulfhydration. This RTS is based on the idea that a reactive cysteine in a protein can be glutathionylated by oxidized glutathione (GSSG) via a thiol/disulfide exchange mechanism. The glutathione-modified cysteine residue can be interchanged to S-sulfhydration by hydrogen sulfide (H_2_S). **(B)** Schematic workflow of the Biotin Switch Technique (BST) to identify glutathionylated protein targets. The proteome in cell extracts was pretreated with TCEP to remove any cysteine-based reversible protein modifications. The reduced proteome was incubated with GSSG (5 mM) to introduce protein S-glutathionylation. After desalting, the GSSG-treated proteome was alkylated, reduced and biotinylated followed by an enrichment with avidin beads. The biotinylated proteins were eluted with TCEP. The proteins being modified by GSSG in the BST eluates can be further analyzed by Western blot assays, or LC-MS. **(C)** Determination of protein S-glutathionylation from mouse liver lysate treated with GSSG. The levels of modified proteins were evaluated in a SDS gel stained with Coomassie blue. The gel lane was excised and digested with trypsin for LC-MS identification. An excess of 2000 protein targets were identified by LC-MS analysis from the liver lysate treated with GSSG. **(D)** Western blot analysis of the BST eluate in C, for two protein targets (MAT1A and GAPDH) being modified by GSSG in mouse liver lysates. The untreated and GSSG-treated liver lysates were subjected to the BST assay. Eluates were analysed by Western blotting for the corresponding proteins.

In order to test our hypotheses, we first evaluated whether the PTM switch from S-glutathionylation to S-sulfhydration could occur on the scale of the cellular proteome. It has been reported that protein S-glutathionylation can be promoted by treatment with oxidized glutathione (GSSG) *in vitro* (Mieyal et al., 2008; Zaffagnini et al., 2012). We therefore incubated protein lysates isolated from mouse livers with TCEP in order to remove reversible cysteine modifications in the proteome. Subsequently, we treated lysates with GSSG to introduce S-glutathionylation. Of note, GSSG-treatment may also result in disulfide bond formation via the initial S-glutathionylation (Mieyal et al., 2008). The GSSG-treated lysates were subsequently subjected to an assay (Anastasiou et al., 2011) referred as the Biotin Switch Technique (BST) to identify the proteins targeted by S-glutathionylation (Fig 5B). We found that the levels of glutathionylated proteins increased by the GSSG treatment (Fig 5C, Supplemental Fig 5), consistent with previous reports (Zaffagnini et al., 2012). MS analysis identified more than 2300 protein targets for S-glutathionylation, including GAPDH (Fig 5C), which supports our previous observation of S-glutathionylation of GAPDH by GSSG treatment (Gao et al., 2015). Western blot analysis confirmed GAPDH and as an additional control, MAT1A (methionine adenosyltransferase 1A) as protein targets being modified by GSSG *in vitro* (Fig 5D).

Does exposure of cell extracts to H_2_S, can convert the proteome-wide S-glutathionylation to S-sulfhydration and what proteins respond to this PTM switch? To address this question, lysates isolated from mouse livers were exposed to TCEP followed by GSSG treatment. After desalting to remove excess GSSG, protein free cysteine thiol groups in the lysates were alkylated with NEM (N-ethylmaleimide) to block remaining free thiol groups. The lysates were incubated with NaHS, then subjected to the BTA assay which was further analyzed by SDS-PAGE (Fig 6A). As shown in Fig 6B, NaHS treatment increased the levels of S-sulfhydrated proteins from the GSSG-treated proteome compared to the untreated control; a dose-dependent increase in the levels of sulfhydrated proteins by H_2_S was also observed (Fig 6C). These data provide strong support for the notion that the PTM switch from S-glutathionylation to S-sulfhydration occurs on a proteome-wide scale. We named this protein cysteine PTM inter-conversion as Redox Thiol Switch S-Glutathionylation to S-Sulfhydration (RTS^GS^).

**Figure 6.**
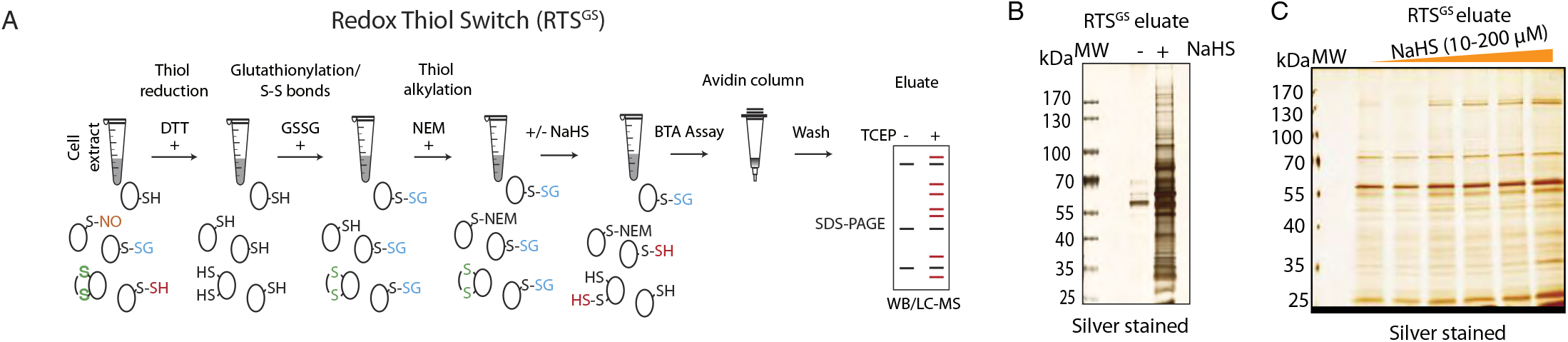
Identification of RTS^GS^ targets by the BTA assay *in vitro.* **(A)** Schematic of workflow to identify cysteine residues within the proteome, candidates for regulation by the RTS^GS^. Cell lysates were pretreated with TCEP to remove any reversible cysteine-based protein modifications, then incubated with GSSG to incorporate S-glutathionylation and disulfides to proteomes. The GSSG-treated protein extracts were treated with N-ethylmaleimide (NEM) to block any free thiol groups. After desalting, the lysates were exposed to H_2_S (NaHS) followed by the BTA assay. The RTS targets were analyzed in a SDS gel or by LC-MS. **(B)** The BTA assay was performed to detect RTS targets in extracts from mouse liver. GSSG-treated mouse liver lysates were divided into two equal fractions, then treated without or with an H_2_S donor (NaHS, 2 mM) for 1h followed by the BTA assay. The eluates were analyzed by SDS–PAGE electrophoresis and silver staining of proteins. **(C)** Mouse liver lysates (equal amount/treatment) were treated and analysed as in B but with increasing concentrations of the H_2_S donor-NaHS (10, 20, 40, 80, 100, and 200 μM).

Pancreatic beta cells are highly sensitive to oxidative stress and the redox status of intracellular glutathione (GSH/GSSG) is important for their metabolism and glucose-stimulated insulin secretion (GSIS) (Gerber and Rutter, 2017). We have previously shown that H_2_S-treatment of MIN6 cells pre-exposed to oxidative stress can reverse the stress-induced inhibitory effects on glucose metabolism (Gao et al., 2015). Therefore, MIN6 cells appeared a good experimental system to test the existence of the RTS^GS^ in living cells and its significance on the cellular metabolism. We hypothesized that small changes in intracellular H_2_S levels can be sensed by metabolic enzymes via the RTS^GS^ mechanism and thus adjust their glucose flux under oxidative stress (Fig 7A, top panel). We therefore tested this concept, by exposing INS1 cells to a protein S-glutathionylation-promoting oxidant for a short time and measuring the effects of H_2_S on the oxidant-induced changes of glucose flux. Diamide is a commonly used oxidant, whose addition to culture medium for short time induces oxidation of glutathione and increases protein S-glutathionylation (Fratelli et al., 2002). Treatment of INS1 cells with diamide for 1 hour, increased the GSSG levels without any global changes in protein synthesis rates (Supplemental Fig 7-1A,B). Diamide-treated INS1 cells were washed to remove diamde, then exposed to NaHS (10 μM) in the presence of glucose [U-^13^C_6_] (Fig 7A, bottom panel). As we previously reported treatment of S-glutathionylated GAPDH with NaHS at concentrations 10-50 μM reversed the inhibition of glutathionylation on the GAPDH activity (Gao et al., 2015). Therefore we considered the concentration of 10 μM of NaHS within the physiological range for the regulation of metabolic activities of enzymes, *in vivo.* As expected, diamide-treatment alone significantly reduced glycolytic and mitochondrial TCA flux as shown by the low ^13^C label incorporation in glyceraldehyde 3-phosphate (G3P), alanine, lactate, citrate, glutamate, succinate, fumarate, malate, oxaloacetic acid (OAA) and aspartic acid (Fig 7A, bottom panel). This inhibition of the metabolic flux was correlated with increased levels of oxidized metabolic enzymes including GAPDH and PCK2 determined by the BST assay (Fig 7B). When the diamide-treated cells were exposed to NaHS, the glycolytic and mitochondrial TCA flux were restored, as shown by the increase in ^13^C-labeling of G3P, alanine, lactate, citrate, glutamate, succinate, malate, OAA. These data suggest that the H_2_S-mediated RTS^GS^ mechanism sustains glycolytic and mitochondrial metabolism under oxidative stress.

**Figure 7.**
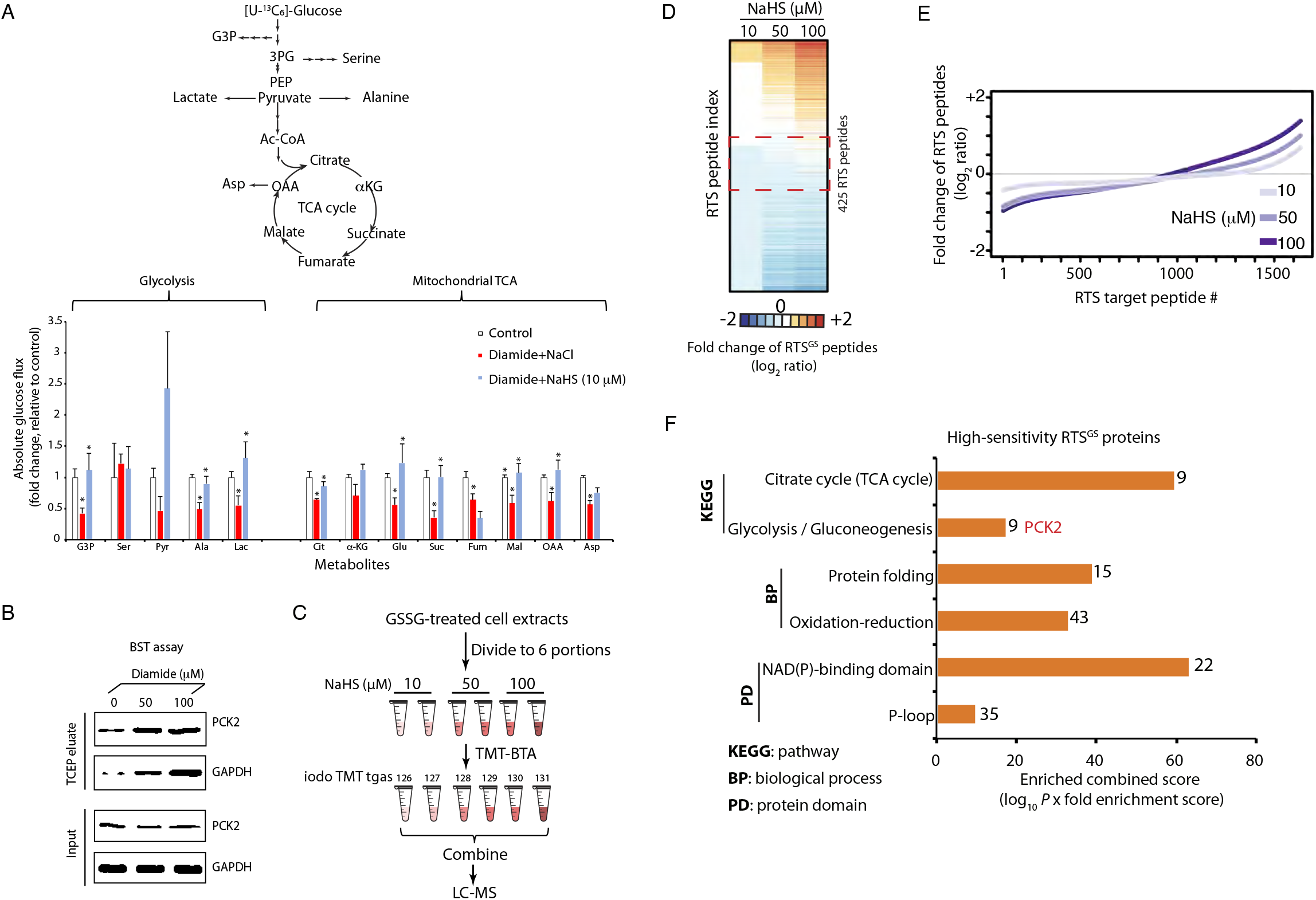
H_2_S restores oxidative stress-decreased metabolic flux in glycolysis and the mitochondrial TCA cycle. **(A)** Measurement of [U-^13^C]-glucose flux in metabolites, expressed as the fold change of absolute glucose flux (ratio of control / treated). INS1 cells were treated with diamide (0.1 mM) in glucose-free growth medium for 90 min, then the cells were switched to 16.8 mM glucose in KRB buffer containing NaHS (0, 10 μM) for 90 min. [U-^13^C]-glucose replaced glucose in the KRB media for the last 90 min of treatments. 4 replicates of each condition were used. **(B)** Evaluation of GAPDH and PCK2 S-glutathionylation in INS1 cells treated with diamide (0, 50 and 100 μM) at the indicated concentrations by the BST assay. INS1 cells were incubated without or with diamide for 90 min, then subjected to the BST assay to determine the levels of glutathionylated GAPDH and PCK2 proteins using western blot analysis (the details of experimental approach for the BST assay was described in Fig 5B). **(C)** Schematic workflow to identify and quantify RTS^GS^ targets by the TMT-BTA assay. MIN6 cell lysates were pretreated with TCEP, then incubated with GSSG. The GSSG-treated lysates were exposed to the H_2_S donor at the indicated concentrations (10, 50 and 100 μM, NaHS), then subjected to the TMT-BTA assay. The RTS^GS^ targets were identified and quantified by LC-MS. **(D)** Heat map showing the relative changes of protein S-sulfhydration in RTS^GS^-target peptides identified by the TMT-BTA assay. 425 RTS^GS^ peptides containing cysteine residues (within the red dotted line frame) displayed higher sensitivity to H_2_S, because a lower concentration of H_2_S (10 μM, NaHS) fully converted S-glutathionylation to S-sulfhydration. **(E)** Curve plot showing the dose-dependent increase of S-sulfhydration levels on RTS^GS^ peptides in the proteome treated with the H_2_S donor from the experimental data in (D). **(F)** Gene ontology analysis of the RTS^GS^ targets in (D), (within the red dotted line frame) showing that among the highly sensitive targets for RTS^GS^, are metabolic enzymes containing a redox domain with a P-loop motif. One of the identified RTS^GS^ targets was the mitochondrial phosphoenolpyruvate carboxykinase 2 (PCK2). Numbers on top of the bars indicate the number of proteins/category.

Based on data in Fig 7A, the concentration of 10 μM NaHS was able to reverse the inhibitory effects of diamide in glucose metabolism. We therefore considered the sulfhydromes formed at 10 μM NaHS treatment as the physiological cellular response to the RTS^GS^ mechanism of regulation. We also hypothesized that the increasing concentrations of NaHS will affect the physiological RTS^GS^ protein sulfhydrome in a manner dependent on the sensitivity of the redox cysteine residues to the RTS^GS^ PTM. We tested this hypothesis in lysates isolated from MIN6 cells. As previously mentioned, the identification of the RTS^GS^ target proteins depends on the sequential conversion of the proteome from S-glutathionylation to S-sulfhydration. Lysates from MIN6 cells were exposed to TCEP and GSSG, followed by the incubation with NaHS at different concentrations 10, 50 and 100 μM, then subjected to the TMT-BTA assay for LC-MS analysis (Fig 7C,D). Quantification of the sulfhydromes between the 10 and 100 μM NaHS treatment, revealed 990 RTS^GS^ protein targets corresponding to 1535 cysteine-containing peptides. About 18% of these peptides exhibited substantial increases (1.5-fold, log_2_ ratio) in S-sulfhydration (Fig 7E), validating the results observed in the SDS acrylamide gel (Fig 6C). Interestingly, we observed that 8% (124 peptides) of the total identified RTS^GS^ cysteine peptides exhibited substantially decreased (1.5-fold, log_2_ ratio) S-sulfhydration, and at least 21 peptides (~17%) contained two cysteine residues in their amino acid sequences. By contrast, only 5 dithiol peptides (~1.8%) were found in the group of RTS^GS^ targets (277 peptides) with increased S-sulfhydration. These data suggest that the group of peptides with decreased S-sulfhydration contain a higher percentage of two cysteine-containing peptides, as compared to the increased S-sulfhydration group. Why did we observe this enrichment in the decreased S-sulfhydration group? Our explanation is that two oxidized cysteine residues in a single peptide may have different sensitivity and reactivity to H_2_S. It is therefore possible that at higher concentrations of H_2_S, one cysteine in a glutathionylated protein is switched to a persulfide by H_2_S, but the second cysteine residue may be reduced to a free thiol, resulting in the retention of the peptide on the avidin column and decrease of the peptide in the eluate (Supplemental Fig 7-2). This can explain the enrichment of dithiol peptides in the decreased S-sulfhydration group. Taken together, we conclude that the concentration of H_2_S in cells under oxidative stress, can be a determinant for the function of the RTS^GS^ mechanism in the regulation of energy metabolism.

Among the 1535 RTS^GS^ peptides, 425 peptides did not change in S-sulfhydration levels with increasing concentration of NaHS (Fig 7D, within the red dotted line frame). This finding suggests that a large number of the RTS^GS^ target proteins have redox cysteine residues that exhibit greater sensitivity to H_2_S. Low levels of H_2_S lead to the complete conversion from S-glutathionylation to S-sulfhydration (Supplemental Fig 7-2). We propose that highly sensitive RTS^GS^ proteins may serve as H_2_S sensors that can quickly switch their oxidized cysteine residues to persulfides under a small magnitude changes of intracellular H_2_S levels in response to oxidative stress. Bioinformatics analysis showed that these protein targets contain NAD(P)H binding motif and they are prevalent in the glycolytic and mitochondrial TCA (tricarboxylic acid) cycle pathways (Fig 7F). The data further suggest that the RTS^GS^ may regulate energy metabolism under oxidative stress conditions. One of those targets is the mitochondrial PEPCK (phosphoenolpyruvate carboxykinase) – (PCK2). Most cells express a cytosolic form of PEPCK (PCK1), an enzyme involved in gluconeogenesis (Yang et al., 2009). The function of PCK2 is not well understood (Yang et al., 2009). PCK2 is the only isoform of PEPCK expressed both in human and mouse pancreatic beta cells (Stark et al., 2009). It has been proposed that the function of PCK2 in beta cells is to regulate GSIS via the mitochondrial GTP-mediated PEP synthesis (Stark et al., 2009). The identification of PCK2 as an RTS^GS^ target strongly suggests its involvement in the redox-mediated regulation of energy metabolism and GSIS.

PEPCK proteins display conserved amino acid sequences across species; they contain a catalytic conserved cysteine residue at the active site (Supplemental Fig 8-1A), suggesting that the activity of PEPCK may be regulated by redox modifications. PCK2 was found as an RTS^GS^ target with high sensitivity to H_2_S (Fig 7D). We therefore tested if PCK2 can be glutathionylated by GSSG, and whether H_2_S can convert S-glutathionylation to S-sulfhydration in the PCK2 protein. We expressed and purified human recombinant PCK2 from *E. coli,* and determined if the enzymatic activity of recombinant PCK2 can be altered by GSSG-mediated S-glutathionylation, or H_2_O_2_-mediated cysteine oxidation sulfenic acid (-SOH), sulfinic acid (-SO_2_h) or sulfonic acid (-SO_3_H). As shown in Figure 8A, both treatments reduced PCK2 enzymatic activity *in vitro;* LC-MS analysis confirmed that the catalytic cysteine 306 in human recombinant PCK2 was predominantly modified by glutathione in response to the GSSG treatment (Supplemental Fig 8-1B-D). Treatment of the glutathionylated PCK2 protein with NaHS, restored its activity (Fig 8B). Furthermore, NaHS-treatment reversed PCK2 S-glutathionylation to S-sulfhydration (Fig 8C). It is shown that the levels of glutathionylated PCK2 were decreased, while the levels of S-sulfhydrated PCK2 were increased by NaHS treatment, confirming the proteomics data (Fig 8C and Supplemental Fig 8-1). These data suggest that S-glutathionylation at the conserved (Supplemental Fig 8-1D) catalytic cysteine 306 of PCK2, has an inhibitory effect on its enzymatic activity.

**Figure 8.**
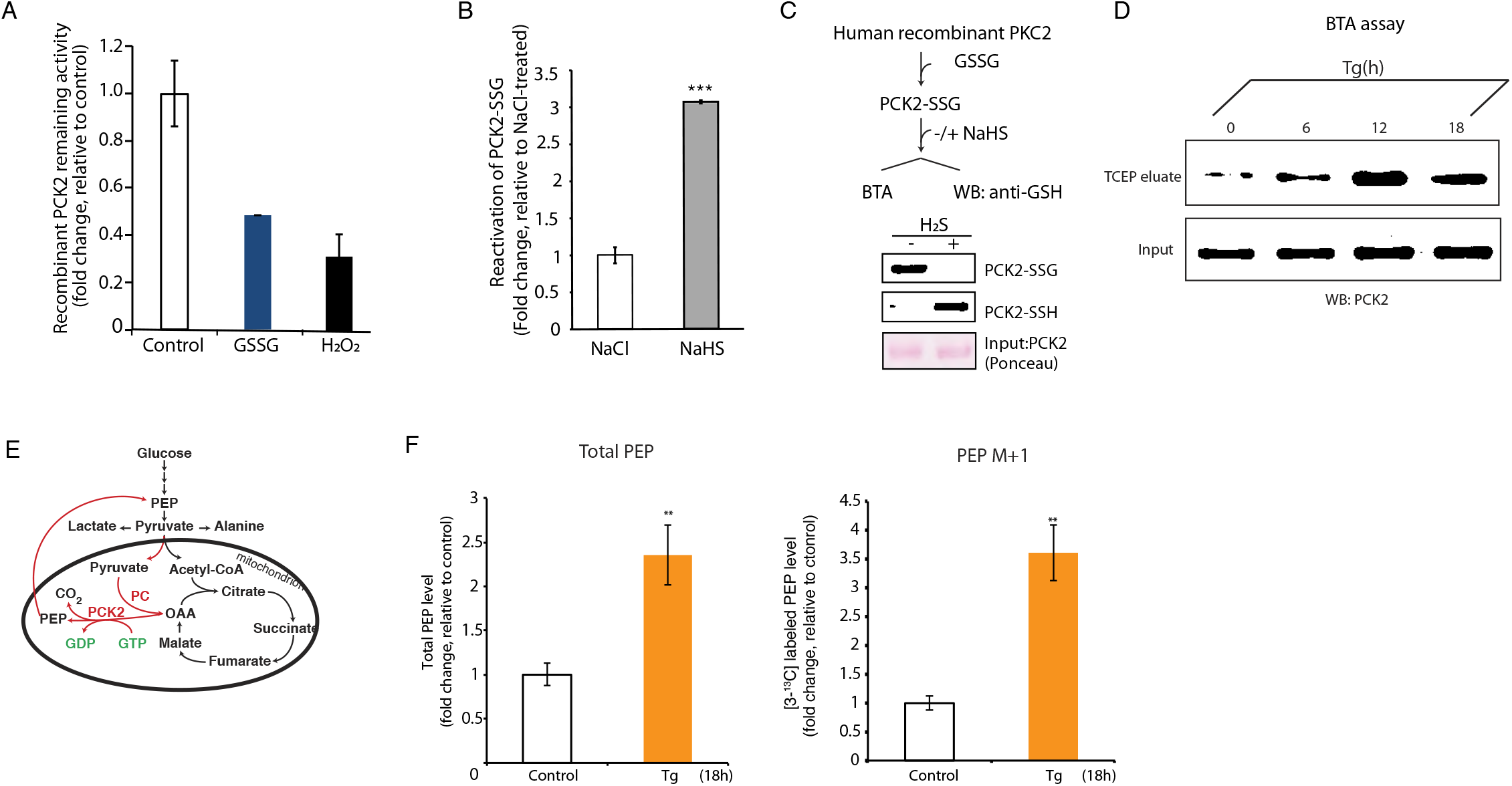
Redox regulation of the PCK2 enzyme by the RTS^GS^ mechanism. **(A)** The enzymatic activity of human recombinant PCK2 is inhibited by GSSG and hydrogen peroxide (H_2_O_2_)-mediated oxidations. Human recombinant PCK2 was treated with GSSG (5 mM) or hydrogen peroxide (1 mM) for 30 min, then subjected to the PCK2 enzymatic activity assay *in vitro.* Triplicates of each condition were used. **(B)** *In vitro* evaluation of the reversal of the inhibitory effect of S-glutathionylation on the activity of the human recombinant PCK2. The recombinant PCK2 was incubated with GSSG for 30 min to introduce S-glutathionylation, then desalted through a gel filtration column. The glutathionylated PCK2 was treated with NaCl as control, or NaHS for 20 min. **(C)** Schematic workflow to determine the RTS^GS^ mechanism in human recombinant PCK2 by Western blot analysis. Human recombinant PCK2 was glutathionylated by GSSG, followed by NaHS-treatment and analysis of the modified protein. Western blot analysis was performed before the BTA assay (lane 1) and after the BTA assay (lane 2) with antibodies against GSH or PCK2 respectively. Glutathionylated PCK2 (PCK2-SSG) levels decrease when exposed to NaHS, while sulfhydrated PCK2 (PCK2-SSH) levels were increased. **(D)** BTA assay identifies PCK2 as a sulfhydrated target in MIN6 cells during ER stress. Western blot analysis of eluates from the BTA assay of sulfhydrated PCK2 in MIN6 cells treated with Tg in a time-dependent manner. A peak of PCK2 sulfhydration occurred at 12h of Tg treatment. **(E)** Diagrammatic representation of phosphoenolpyruvate (PEP) cycling mediated by mitochondrial PCK2. **(F)** Evaluation of the PCK2 activity in MIN6 cells under chronic ER stress by a metabolic labeling approach. MIN6 cells were treated with Tg (Gao et al., 2015) in culture media for 18 h. During the last 3.5 h of Tg treatments, sodium pyruvate [^3^C_13_] replaced unlabeled pyruvate in the growth media. The total levels of PEP and PEP M+1 were determined by LC-MS analysis showing that chronic ER stress increases the enrichment of total PEP and PEP M+1 in MIN6 cells. Four replicates of each condition were used.

We finally tested the sulfhydration status of PCK2 during chronic ER stress, a condition that associates with increased ROS levels, and increased H_2_S synthesis (Gao et al., 2015). We previously showed that chronic ER stress in MIN6 cells promotes protein S-sulfhydration which restores stress-induced inhibition of glycolysis (Gao et al., 2015). MIN6 cells were treated with the chemical stressor thapsigargin (Tg) to induce chronic ER stress (Guan et al., 2017). Treatment with Tg induces the ATF4/CTH axis of protein S-sulfhydration (Gao et al., 2015). After incubation for 6, 12 and 18h, the cells were lysed and subjected to the BTA assay. As expected, the levels of sulfhydrated PCK2 were increased in response to ER stress (Fig 8D). LC-MS analysis confirmed that the catalytic cysteine 306 was sulfhydrated from the immunepurified FLAG-PCK2 protein expressed in Tg-treated MIN6 cells for 18h (Supplemental Fig 8-2A-C). This prompted the question as to whether PCK2-mediated flux of PEP (phosphoenolpyruvate) is regulated by the H_2_S-dependent modification of this enzyme. PCK2 converts OAA to PEP using GTP as a cofactor (Fig 8E). To measure the changes in PCK2-mediated PEP flux, MIN6 cells were treated with Tg for 18h; the growth media was changed to contain the stable isotope [3-^13^C] pyruvate during the last 3.5 h of treatment. Tg-treatment augmented total and M+1 PEP in MIN6 cells (Fig 8F). These data were consistent with our previous observation that H_2_S synthesis during chronic ER stress restores stress-induced inhibition of glycolytic flux (Gao et al., 2015). We conclude that enzymes in energy metabolism sense increased H_2_S levels under conditions of chronic oxidative stress and via the RTS^GS^ mechanism rescue stress-induced deficits in energy metabolism.

## Discussion

The gas hydrogen sulfide (H_2_S), induces S-sulfhydration of redox sensitive cysteine residues in the proteome and thus regulates protein functions. The development of quantitative tools that can profile proteome-wide S-sulfhydration is crucial to identify the major protein targets and determine their relationship with the biological effects of this gas. Here, we introduced the TMT-BTA assay, which allowed the quantitative assessment of changes in protein-cysteine sulfhydration *in vivo* and *in vitro* under various cellular contexts. Using multiple validations, our data confirm the specificity and suitability of the TMT-BTA assay for the analysis of protein S-sulfhydration. With this new quantitative proteomics approach, we have demonstrated a good correlation of the S-sulfhydration status of intracellular proteins with the expression of H_2_S-generating enzymes. This PTM occurs only on redox sensitive cysteine residues of specific proteins. Our findings suggest the protein S-sulfhydration mechanism as a dial switch in the regulation of cellular metabolism in pathophysiological states. Glycolytic proteins are the most highly sulfhydrated proteins in EndoC-BH3 cells, including (ALDOA), an enzyme that converts fructose 1,6 bisphosphate (FBP) into dihydroxyacetone phosphate (DHAP) and glyceraldehyde 3 phosphate (G3P). The regulation of FBP levels via synthesis and degradation is critical to control GSIS (Kebede et al., 2008), the most important function of beta cells. ALDOA exhibits conserved amino acid residues across species, including the two cysteine residues (Cys^73^ and Cys^178^) which were extensively sulfhydrated in human pancreatic beta cells (Fig 3F). A previous study showed that plant cytosolic ALDOA is subject to both S-glutathionylation and S-nitrosylation at the same cysteine residues, both resulting in inhibition of its enzymatic activity (van der Linde et al., 2011). Given this correlation, we speculate that S-sulfhydration of Cys^73^ and or Cys^178^ might regulate ALDOA enzymatic activity, change glycolytic flux, and affect insulin secretion in pancreatic beta cells.

There are three H_2_S-generating enzymes, CTH, CBS and MST, in mammalian cells (Sbodio et al., 2018). Unlike the other two, CTH protein expression is transcriptionally induced by the stress-induced master transcription factor-ATF4, thus linking H_2_S synthesis to diverse physiological and pathological stress conditions (Paul and Snyder, 2015; Sbodio et al., 2018). The ATF4/CTH-mediated changes in the sulfhydrome are shown here for the first time. Pancreatic beta cells require the presence of active stress response mechanisms for their function, due to the easily disturbed proteostasis in the ER, during fluctuations of proinsulin synthesis (Liu et al., 2010). Biochemical studies with recombinant insulin showed that the insulin B chain was a target for the ATF4/CTH-mediated S-sulfhydration, and thus prompted the question on the physiological significance of this regulation. Pancreatic beta cells synthesize proinsulin on ER-associated ribosomes. Proinsulin is then folded and processed to produce insulin which is stored in secretory granules and is secreted in response to nutrients, such as glucose (Liu et al., 2010). Increased H_2_S levels repressed GSIS in INS1 cells treated with H_2_S donors (Supplemental Fig 9) (Yang et al., 2007). The mechanism via which H_2_S decreases GSIS is not well understood. It has been shown that S-sulfhydration of the K_ATP_ channel increased the activity of the ion channels, resulting in an inhibition of GSIS in pancreatic beta cells (Yang et al., 2005). We have shown here that in addition to this mechanism, S-sulfhydration of insulin (specifically the B chain) may also contribute to inhibition of GSIS. Insulin B chain cysteine sulfhydration is expected to break the disulfide bonds and unfold its structure (Fig 4K), reducing the folded insulin in the secretory granules. This may consist a novel physiological mechanism for the fine-tuning of GSIS in pancreatic beta cells. Impairment of GSIS in pancreatic beta cells is associated with the development of diabetes in humans. A recent study showed that the insulin B chain fragment triggered an activation of immune T cells by enhancing the pathogenic B chain peptide loading to antigen-presenting cells, which contributed to the development of type 1 diabetes (Wan et al., 2018). Animal and cell culture models have showed that elevated H_2_S production is associated with the development of diabetes (Szabo, 2012). Indeed, whether S-sulfhydration of the insulin B chain in islets is an onset to trigger T cell activation leading to beta cell destruction remains to be further investigated.

We discovered that a large number of protein targets (~990 targets) were subjected to the RTS^GS^ PTM switch from S-glutathionylation to S-sulfhydration mediated by hydrogen sulfide. A key feature of this PTM switch is that both modifications occur on the same cysteine residue. The RedoxDB database, indicated about 9% of the identified RTS^GS^ peptides are known to be modified by cysteine-based PTMs including disulfides, S-nitrosylation, S-glutathionylation and sulfenic acid. This is in agreement to our functional annotation analysis that RTS^GS^ proteins contain redox sensitive cysteine residues which are highly susceptible to oxidative stress associated PTMs. Pathway analyses also showed that RTS^GS^ targets with the highest sensitivity to H_2_S were involved in energy metabolism. We can therefore propose that the RTS^GS^ mechanism may regulate energy metabolism under chronic oxidative stress. Five RTS^GS^ targets involved in glycolysis and the mitochondrial TCA cycle were found: Glyceraldehyde 3-phosphate dehydrogenase (GAPDH), pyruvate kinase 2 (PKM2), isocitrate dehydrogenase (IDH), oxoglutarate dehydrogenase (OGDH) and pyruvate dehydrogenase (PDH). The enzymatic activities of those enzymes are inhibited by redox modifications under increased ROS levels (Mailloux et al., 2014). Thus, H_2_S-mediated RTS^GS^ of these metabolic enzymes is likely part of the mechanism to reverse the ROS-induced inhibition of glycolysis and the TCA cycle and restore metabolic flux under oxidative stress conditions.

One novel RTS^GS^ target was the mitochondrial enzyme PCK2. This enzyme converts OAA to PEP and promotes insulin secretion in pancreatic beta cells (Stark et al., 2009). Both isoforms of PEPCK (PCK1 and PCK2) displayed a conserved catalytic cysteine at their active site (Supplemental Fig 7-1D). In human and rodent pancreatic beta cells, only the mitochondrial PCK2 enzyme is expressed, whereas other tissues express both PCK2 and PCK1. It is known that oxidative stress is a hallmark of ER stress (Cao and Kaufman, 2014). PCK2 is a redox sensitive enzyme regulated by the RTS^GS^ described in the current study. This regulation mediated by H_2_S might indicate a protective mechanism to sustain mitochondrial function and restore insulin secretion in pancreatic beta cells under chronic ER stress. With higher ROS production in response to ER stress, mitochondrial oxidative phosphorylation and TCA metabolic rates are decreased to preserve mitochondria (Gao et al., 2015). The oxidation of the PCK2 enzyme results in an accumulation of OAA within the mitochondria due to the reduction of PEP cycling (Gao et al., 2015). The restoration of PCK2 activity triggered by increased H_2_S under chronic ER stress could give beta cells the capability to increase ATP turnover and normalize the redox state. This in turns helps to reduce oxidative stress imposed on the cells. We therefore view that the H_2_S-mediated RTS^GS^ switch in mitochondrial PCK2 is an adaptive mechanism for promoting beta cell survival, along with the improved function of pancreatic beta cells.

The H_2_S-mediated protective effect against oxidative stress-mediated-deregulations of biological activities, is an evolutionary conserved process (Filipovic et al., 2018). For example, endogenous H_2_S production in multiple bacterial species offers resistance to oxidative stress and various classes of antibiotics (Mironov et al., 2017). H_2_S reverses the age-associated impairment of the formation of new blood vessels in muscle (Das et al., 2018), and the increase in H_2_S production in endothelial cells induced by sulfur amino acid restriction promotes glucose uptake and ATP production via glycolysis (Longchamp et al., 2018). However, the protective effects of H_2_S on cellular metabolism, may also enhance tumor growth, which is not a desired outcome for cancer therapeutics. Our studies can transform our current thinking of cancer cell metabolism. Tumors exhibit increased ATP production through glycolysis (Vander Heiden et al., 2009) and high H_2_S levels in tumors are associated with tumor growth (Szabo et al., 2013). In addition, unique splicing isoforms of key metabolic enzymes contribute to increased glycolytic flux in tumors (Luo and Semenza, 2012). For example, tumors express an alternatively spliced form of pyruvate kinase, PKM2, which enhances aerobic glycolysis and provides tumor growth advantage (Anastasiou et al., 2011). PKM2 is a redox sensitive protein with cysteine oxidation being inhibitory of its activity. Given that we have identified PKM2 as an RTS^GS^ target, we propose that the H_2_S-mediated PTM switch on metabolic enzymes such as PKM2, is a potential mechanism for tumor growth.

In the current study, we have developed an experimental tool, named the TMT-BTA assay, for selectively trapping and tagging cysteine persulfides in the proteome of cells. Combined with click chemistry and quantitative MS proteomics, the TMT-BTA assay revealed the natural protein sulfhydrome of two human cell lines, and identified insulin as a novel sulfhydration substrate in pancreatic beta cells. With this approach, we have identified biological pathways being targets for RTS^GS^, a specific reversible thiol modification in cysteine residues under oxidative stress conditions. The findings suggest that the levels of H_2_S-producing enzymes control the sulfhydromes of cells, thus modulating energy metabolism. The identified pool of protein targets subjected to the RTS^GS^ can serve as a valuable resource to study the effects of H_2_S on redox homeostasis and metabolism in physiological and pathological states. Although in physiological states, H_2_S may protect normal cells from ROS-induced metabolic deregulation, in cancer cells, it may be an unwanted passenger. Cancer cells are known to reroute signaling pathways to gain a metabolic advantage to prevent oxidative damage under stress conditions, which is a signature phenotype associated with most tumors (Vander Heiden et al., 2009). We therefore view that the H_2_S-mediated RTS^GS^ may be a major contributing mechanism to reprogram cancer cell metabolism for their proliferation and survival. In this case, it would provide a rationale for cancer therapeutics to modulate the activity of the H_2_S-producing enzymes in tumor cells.

## Acknowledgements

We thank Dr. Ruma Banerjee for providing recombinant CTH used in this study. We thank Dr. Danny Manor for providing IHH cells used in this study. We also thank Dr. Renliang Zhang for assisting with advice on the PEP flux analysis. We thank Ken Farabough for provided editorial assistance. We acknowledge the Univercell Biosolutions (Paris, France) providing the EndoC-BH3 cells used in this study. This work was supported by NIH grants R37-DK060596, R01-DK053307 (to MH), American Diabetes Association Postdoctoral Fellowship 1-16-PDF-018 (to X.-H. G). The Orbitrap Elite and Fusion Lumos LC-MS instruments were purchased via an NIH shared instrument grants 1S10RR031537-01, 1S10OD023436-01 (to BW).

## Author Contributions

Xing-Huang Gao and Maria Hatzoglou conceived the study; designed and performed the experiments; wrote the manuscript and analysed and interpreted the data. Ling Li and Belinda Willard, performed the proteomics analysis with LC-MS. Marc Parisien and Luda Diatchenko, performed bioinformatics tasks and assisted in data interpretation.Matt Mcleod, expressed and purify human recombinant PCK2 protein.Jing Wu, performed Western blot analysis.Ilya Bederman, performed and analysed the metabolic flux studies. Zhaofeng Gao, performed bioinformatics tasks and assisted in data interpretation and manuscript editing. Dawid Krokowski, assisted in data interpretation and manuscript editing. Steven M Chirieleison and Derek Abbott, contributed the Crispr-Cas9 experiments. Vivien Yee, adviced on structural modeling of PCK2. Charles L. Hoppel, provided expertise on cellular metabolism, mitochondrial function and metabolic data analysis. Richard G Kibbey, provided expertise on the PEP flux study.Todd Holyoak, advised on biochemical characterization of the human recombinant PCK2. Peter Arvan, provided expertise on beta cell biology and glucose stimulated insulin secretion assays.

---

**Supplementary Figure 1.**
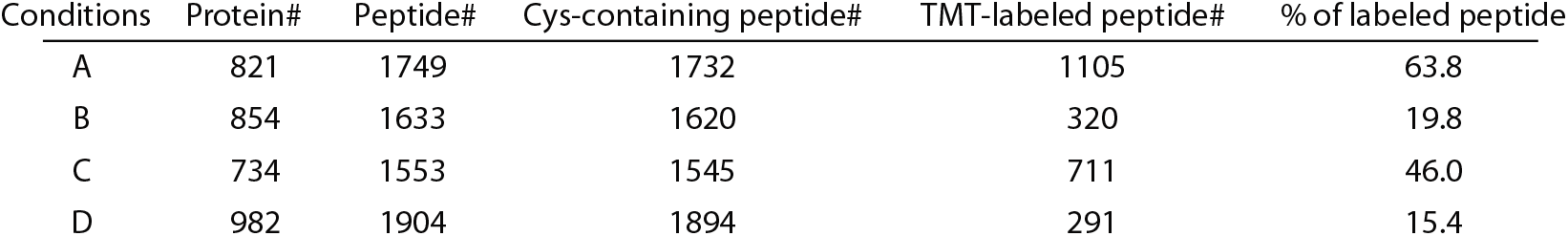
Optimization of the reaction buffer used for sulfhydrated peptide labeling by the iodoTMT tags. Cell extracts treated NaHS (100 μM) were subjected to the TMT-BTA assay. After biotinylation and trypsin digestion, the sample was divided into four equal fractions. Each fraction was incubated with the same amount of avidin beads, the biotinylated peptides were eluted with the different buffers as described below: A. 50 mM TEAB (triethylamonium bicarbonat) and 10 mM TCEP, pH 8.0 B. 50 mM HEPES-NaOH and 10 mM TCEP pH 8.0 C. 50 mM TEAB and 20 mM TCEP, pH 8.0 D. 50 mM HEPES-NaOH and 20 mM TCEP, pH8.0 The BST eluates were mixed with iodoTMT reagents and incubated for 1h. The reactions were stopped by adding DTT (20 mM). After desalting to remove excess chemicals. All the samples were combined for LC-MS identification and quantification.

**Supplementary Figure 3.**
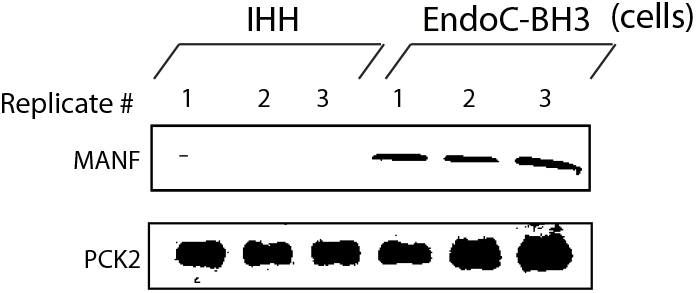
Assessment of protein levels in IHH and EndoC-BH3 cells. Western blot analysis for the indicated proteins, of cell extracts isolated from IHH and EndoC-BH3 cells.

**Supplementary Figure 4-1.**
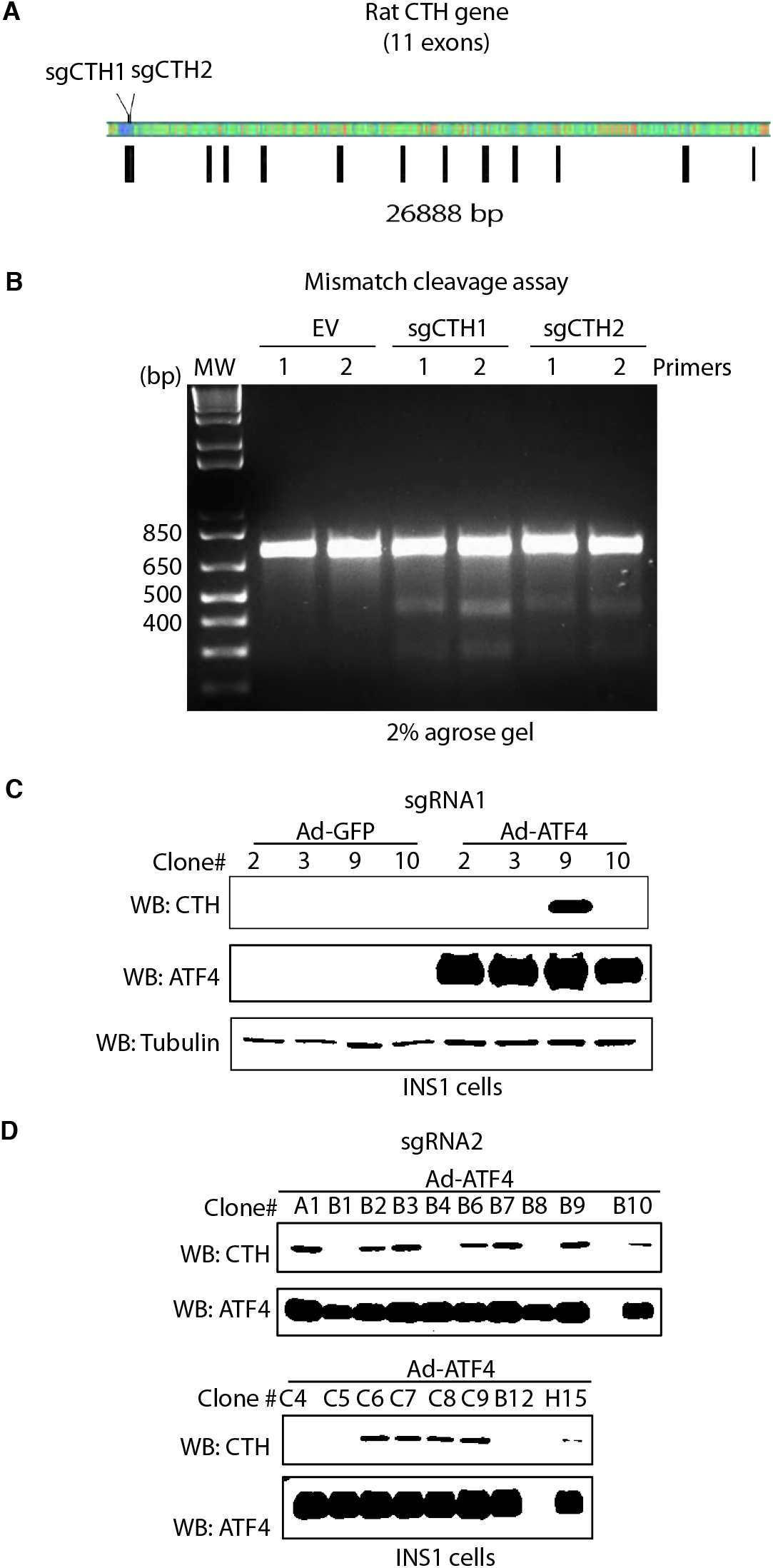
Assessment of CRISPR–Cas9-mediated CTH gene knockout in INS1, rat pancreatic beta cells. **(A)** Schematic representation of the rat CTH gene *loci* and two guide RNA targeting sites. The binding sites by the guide RNAs were overlapped with the translation initiation codon in exon1. **(B)** Evaluation of CTH gene knockout by the mismatch detection assay. INS1 cells were transduced with lentivirus to express the indicated sgCTH RNAs, or empty vector (EV) followed by selection with puromycin to enrich for CTH knockouts. Two different PCR primers (set 1 and set 2) were used to amplify the regions flanking the CRISPR targeting sites. Insertion and deletions in these genomic PCR products were detected by the mismatch assay. **(C, D)** Western blot analysis of INS1 cell clones CRISPR-CTH overexpressing ATF4. Individual INS1 cell clones were transfected to express ATF4 for 48h, then lysed. The levels of CTH protein in each clone were evaluated by Western blot analysis.

**Supplementary Figure 4-2.**
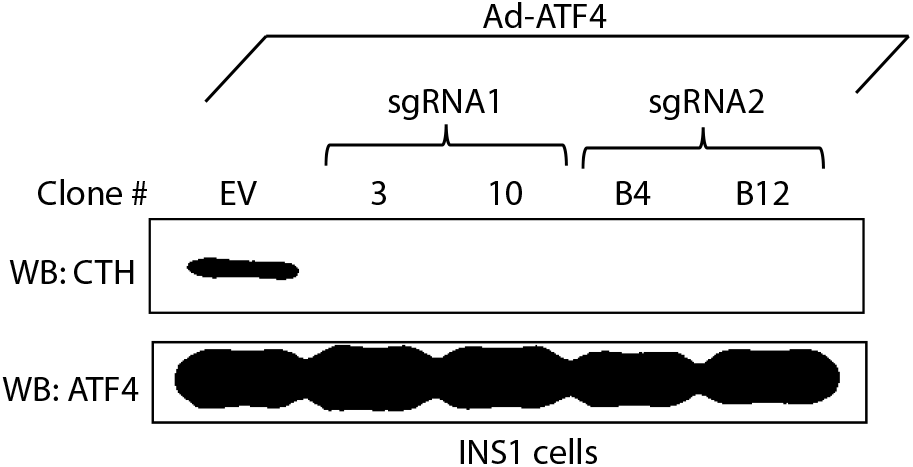
Verification of CRISPR-cas9 mediated CTH gene knockout in INS1 cells expressing ATF4. Individual clones derived from two different guide RNAs were transfected to express ATF4. The levels of ATF4 protein were evaluated by Western blot analysis.

**Supplementary Figure 5.**
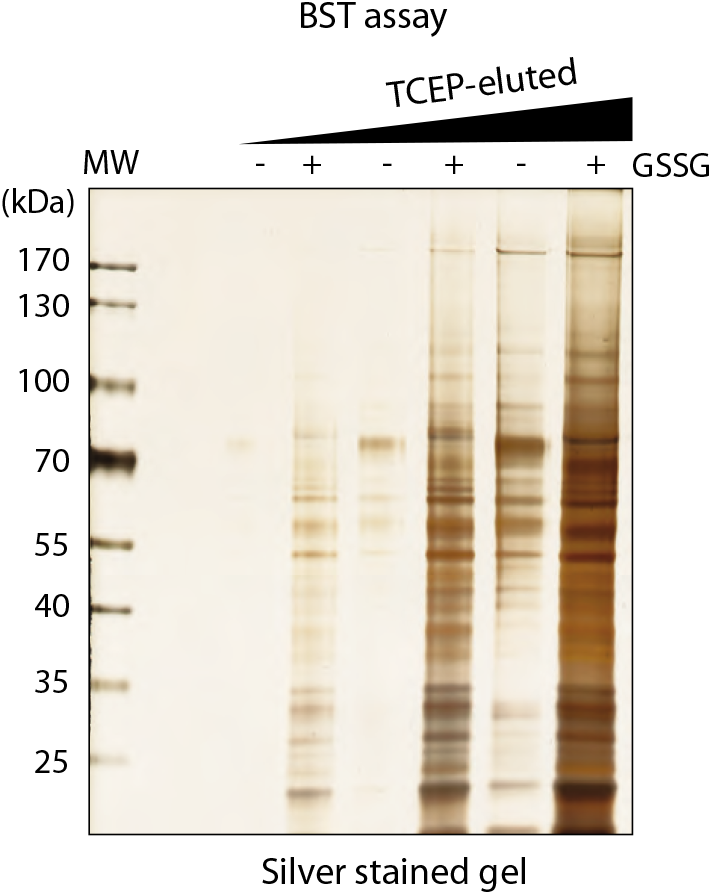
Detection of glutathionylated proteins by the Biotin Switch Technique (BST). Mouse liver lysates were pretreated with TCEP (20 mM) to remove any reversible cysteine protein modifications. After desalting, the lysates were exposed to GSSG (1 mM) to introduce S-glutathionylation, followed by the BST assay. After biotinylation of GSSG-modified proteins, the lysates were divided into six equal fractions, were bound on streptavidin columns and eluted by the addition of increasing concentrations of TCEP (1-20 mM). The eluates were analyzed by SDS–PAGE electrophoresis and silver stained.

**Supplementary Figure 7-1.**
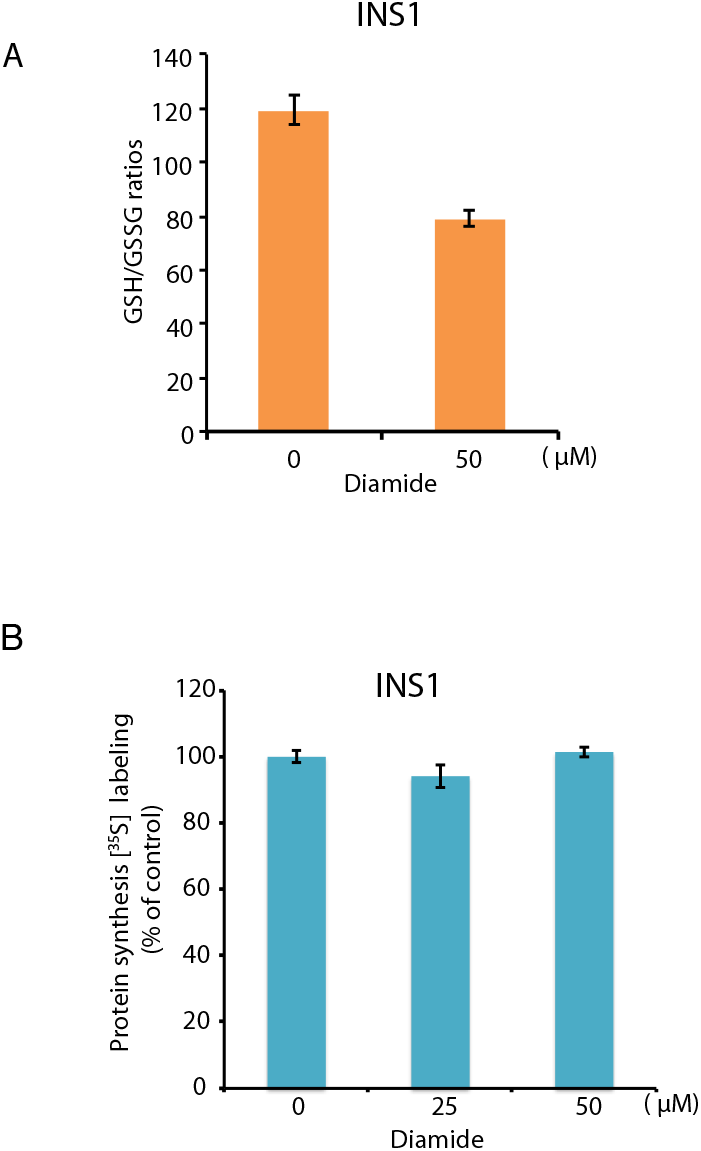
**(A)** Diamide treatment increases oxidation of intracellular glutathione (GSH). INS1 cells were exposed with diamide at the indicated concentrations for 1h. The intracellular GSH/GSSG ratios were quantified by the GSH/GSSG-Glo^TM^ assay. Results are the mean of triplicate determinations. **(B)** Protein synthesis rate is unaffected by diamide treatment. Protein synthesis was measured by [^35^S] Met/Cys incorporation into proteins in INS1 cells treated with diamide at the indicated concentrations for 1h. Results are the mean of triplicate determinations.

**Supplementary Figure 7-2.**
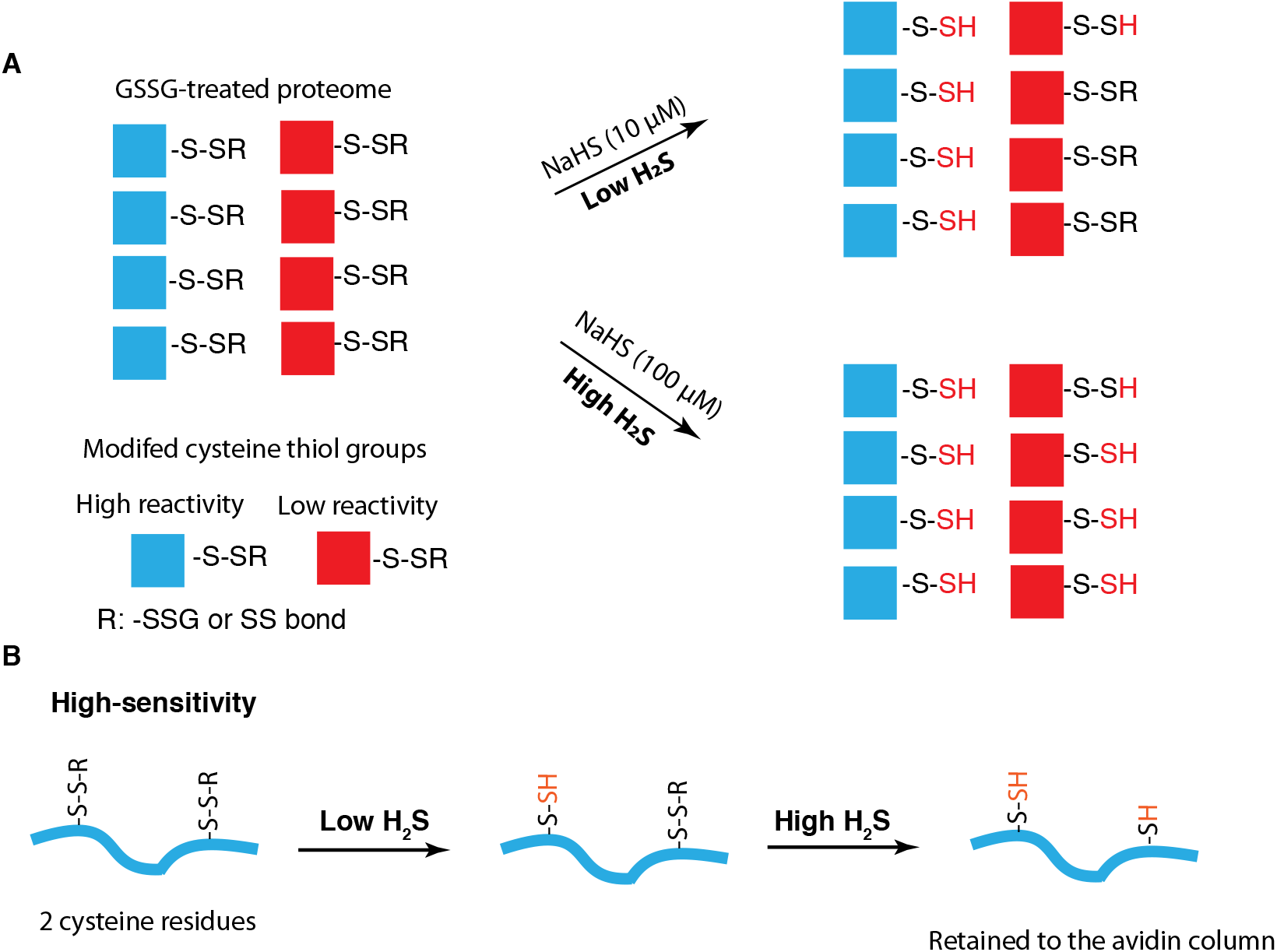
Schematic representation of the predicted RTS^GS^ peptides in the eluate of the TMT-BTA assay as a consequence of increasing concentrations of H_2_S. **(A)** At low concentrations (10 μM), the modified cysteine residues including disulfides and S-glutathionylation are discriminated for S-sulfhydration *(top).* The modified cysteine residues with lower reactivity (red) should show a concentration-dependent increases in S-sulfhydration *(bottom).* **(B)** Model of a high sensitivity peptide containing two modified cysteine residues, one that can be sulfhydrated at low H_2_S concentrations, and a second cysteine residue that is reduced to a free SH group only at high concentrations (100 μM), leading to the peptide retained to the streptavidin beads. This differential response of close proximity cysteine residues, can explain the decreased S-sulfhydration observed for some proteins with the TMT-BTA assay.

**Supplementary Figure 8-1.**
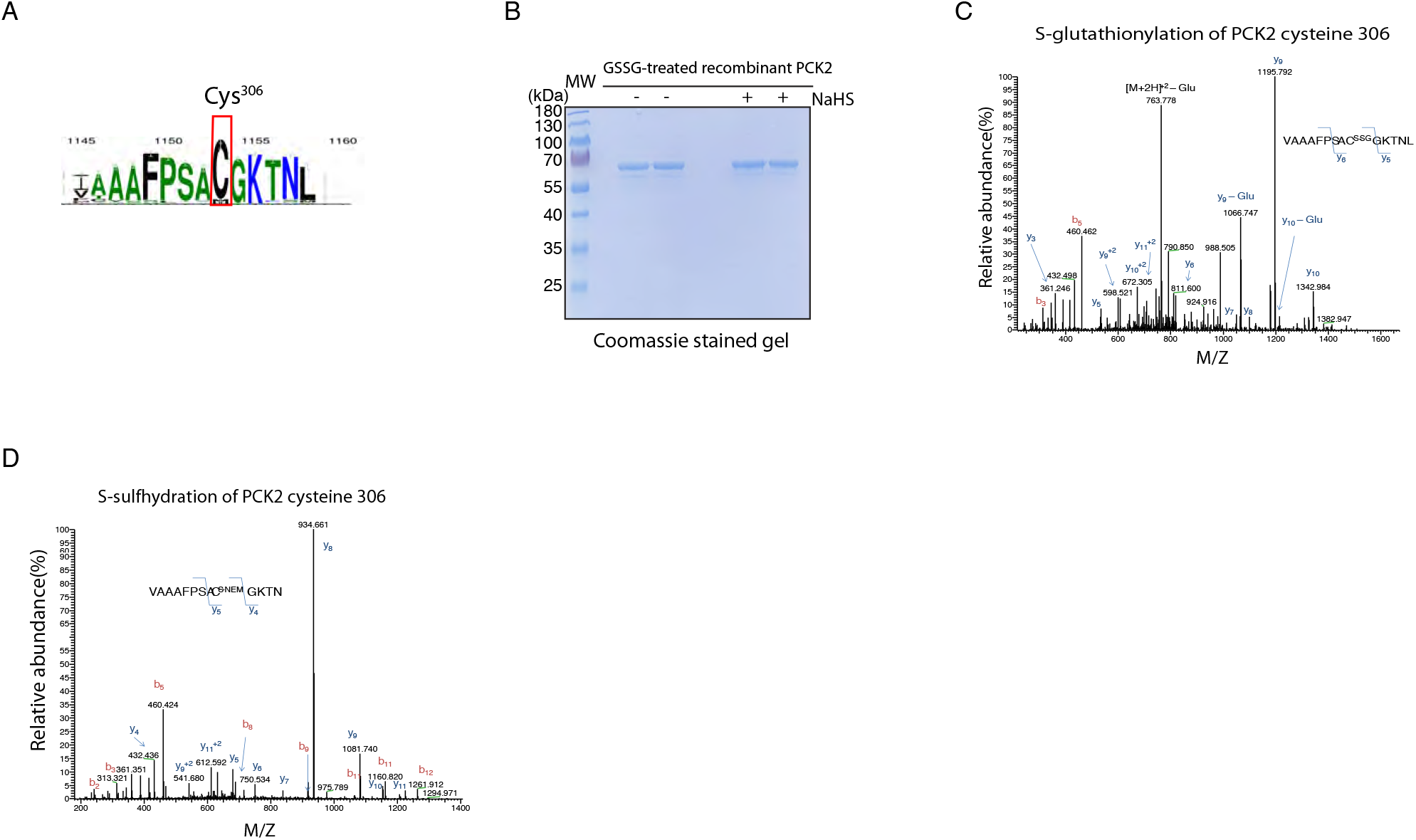
**(A)** PEPCK protein sequence alignment showing the catalytic cysteine 306 position. LOGO representing the cysteine^306^ residue-containing sequence of the PEPCK catalytic domains in different species. The overall height of each letter indicates the degree of amino acid sequence conservation. **(B)** LC-MS analysis identifies S-glutathionylation and S-sulfhydration of human recombinant PCK2 at the catalytic cysteine, Cys^306^. Human recombinant PCK2 was treated with GSSG (5 mM) to introduce S-glutathionylation, then incubated with NaHS to induce S-sulfhydration. After desalting, both GSSG and GSSG/NaHS-treated PCK2 proteins were alkylated by NEM to block any free SH groups. The samples were resolved by SDS–PAGE electrophoresis followed by a LC-MS analysis. **(C)** GSSG-treated PCK2 shows a mass shift of Cys^306^ corresponding to a glutathione molecule. **(D)** NaHS-treated PCK2-SSG shows a mass shift of Cys^306^ corresponding to a NEM conjugated-persulfide.

**Supplementary Figure 8-2.**
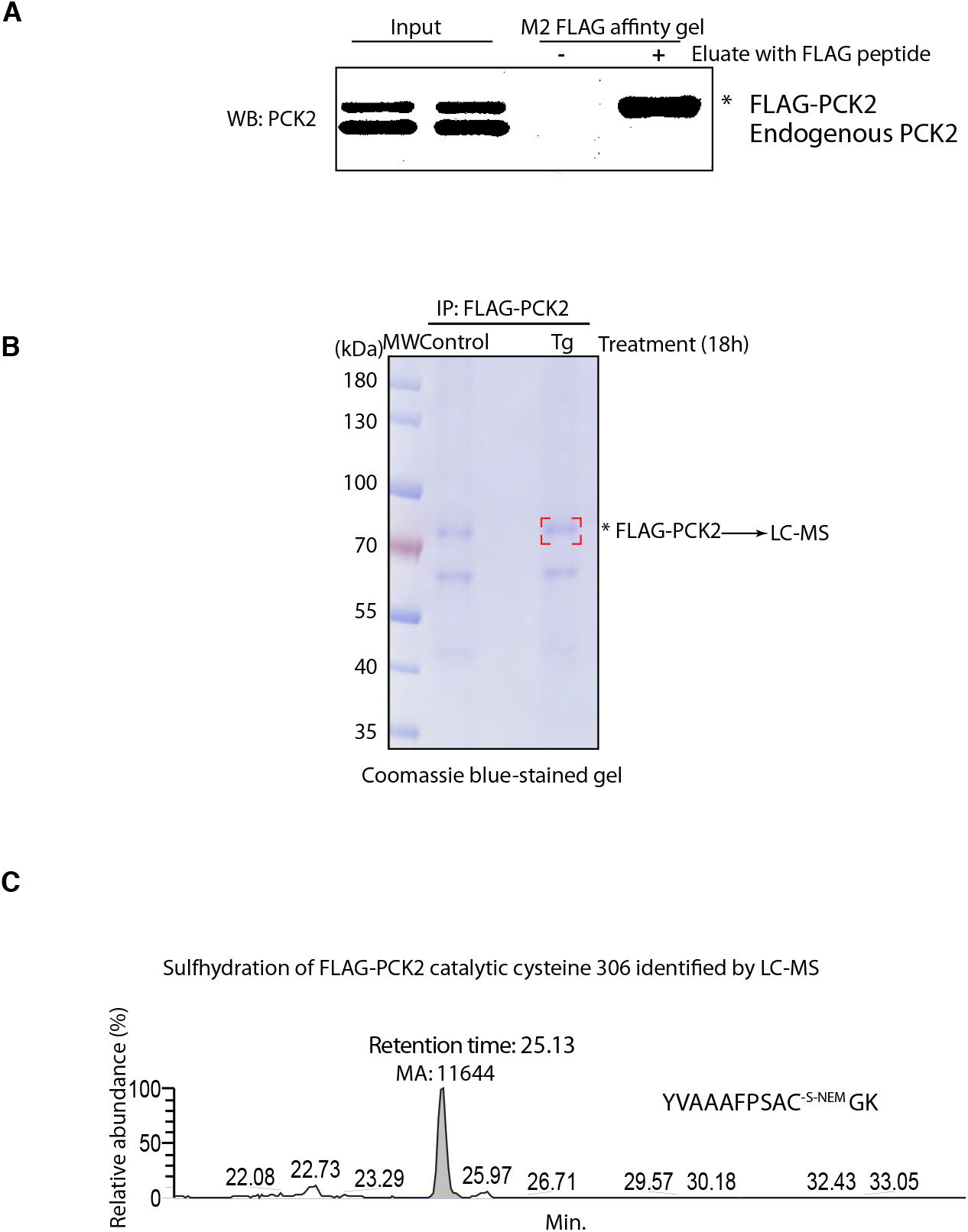
Chronic ER stress promotes S-sulfhydration of FLAG-PCK2 at the catalytic cysteine 306 in mouse pancreatic beta cells. **(A)** Immunoprecipitation of human FLAG-tagged PCK2 expressed in MIN6 cells. MIN6 cells were transfected to express human FLAG-tagged PCK2. After 48h of transfection, the cells were lysed, and subjected to immunoprecipitation. The FLAG-PCK2 was eluted by a FLAG peptide under non-reducing conditions. The levels of FLAG-PCK2 protein were evaluated by western blot analysis. **(B)** MIN6 cells expressing FLAG-PCK2 were treated with Tg for 18h, then subjected to an immunoprecipitiation of the FLAG fusion PCK2. The eluates were resolved by a SDS-gel, the bands were excised and digested for LC-MS analysis. **(C)** LC-MS identifies the immunopurified FLAG-PCK2 catalytic cysteine 306 as being modified by S-sulfhydration in MIN6 cells treated with Tg after 18h.

**Supplementary Figure 9.**
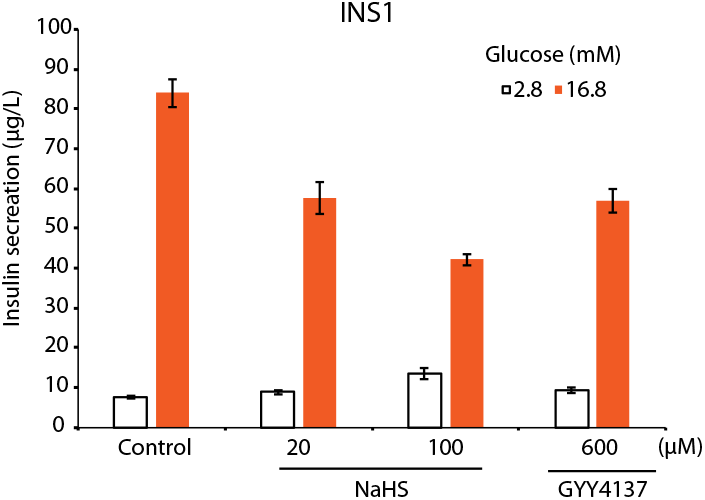
Treatment of INS1 cells with H_2_S donors inhibits glucose-stimulated insulin secretion (GSIS). INS1 cells were treated with two different H_2_S donors (NaHS, GYY4137) at the indicated concentrations for 1h, then subjected to the GSIS assay. The levels of released insulin were quantified by an ELISA assay. 5 replicates of each condition were used.

## STAR Methods

Antibodies

CBS: Abnova, H00000875-001p

CTH: Sigma, HPA023300

ATF4: Santa Cruz Biotechnology, SC-200

GAPDH: Santa Cruz Biotechnology, SC-32233 (6C5)

GSH: Virogen, 101 −A (D8 clone)

MANF: Icosagen AS, 310-100

3MST: Santa Cruz Biotechnologies, SC-376168

### Cell line and cell culture

Human pancreatic beta cells (EndoC-BH3) were purchased from the Univercell Biosolutions (Paris, France). The EndoC-BH3 cells were cultured in DMEM containing 5.6 mM glucose, 2% BSA fraction V, 50 μM 2-mercaptoethanol, 10 mM nicotinamide, 5.5 μg/ml transferrin, 6.7 ng/ml sodium selenite, penicillin (100 units/ml) and streptomycin (100 μg/ml). Ten μg/ml of puromycin (selective antibiotic) were added in the complete medium. The cells were seeded onto matrigel- and fibronectin-coated culture plates at 4 × 10^6^ cells/plate.

MIN6 and INS1 cells were cultured as described previously (Gao et al., 2015).

### Glucose-Stimulated Insulin Release (GSIS)

Glucose-stimulated insulin release was assayed INS1 cells as described previously (Haataja et al., 2013).

### Quantification of the intracellular ratio of oxidized and reduced glutathione

INS1 cells (5×10^4^ cell per well) were seeded into 96-well plates. After 48 h the growth media was removed, the total glutathione contents and GSH/GSSG ratios were determined with the GSH/GSSG-Glo^TM^ Tm assay from Promega.

### Construction, Expression and purification of human recombinant PCK2 protein

The human PCK2 gene was inserted into peSUMOstar Kan vector from Life Sensors as previously described (Johnson and Holyoak, 2012), beginning at Leu33, corresponding with the mature N-terminal residue that would be present after import into the mitochondria and removal of the mitochondrial signal peptide.

Expression of human PCK2 constructs began with four 50mL overnight cultures of LB and kanamycin (50 μg/ml) grown at 37°C. These cultures were spun down and added to four 1L flasks of LB with kanamycin. They were grown at 37°C until an OD_600_ of 1.2 in which they were induced with 0.1 mM IPTG and the temperature was reduced to 20°C. Cells were harvested 20-24 hours later.

Human PCK2 purified as follows: Cell pellets were resuspended in buffer 1 (10 mM KPO_4_^-^ pH 6.8, 10% (w/v) glycerol, 300 mM NaCl, 10 mM imidazole, 2 mM TCEP) and lysed via two passages through a pre-chilled French press pressure cell at 1000 psi. The lysate was spun down at 12,000 g for 1 hour at 4°C (all subsequent steps were completed at 4°C). The supernatant was removed and added to pre-equilibrated with (buffer 1) nickel-NTA resin in which it was stirred for 1 hour (UBP-BIO). The protein-bound resin was washed with buffer 1 until the flow-through had an A_280_ less than 0.1. Protein was eluted with buffer 2 (10 mM KPO_4_^-^ pH 6.8, 300 mM imidazole. 2 mM TCEP) and fractions containing protein were concentrated to 5 ml. The protein was buffer exchanged into buffer 3 (10 mM KPO_4_^-^ pH 6.8, 2 mM TCEP) by passage through a preequilibrated (in buffer 3) P6-DG column (BioRad). This protein was loaded onto CHT type 2 resin (BioRad) via FPLC and washed with buffer 3. The column was washed with a linear gradient (10-400 mM KPO_4_^-^ pH 6.8) and the peaks with PCK activity were collected, concentrated, and buffer exchanged as above into buffer 3. The His-tagged PCK2 protein was then treated overnight with SUMO protease to cleave the N-terminal fusion tag. Cleaved PCK2 was then added to clean, pre-equilibrated in (buffer 3) nickel-NTA resin and stirred for 1 hour. After rebinding, the majority of protein precipitated. The remaining PCK2 was collected as flow through. Fractions containing protein were then concentrated, and buffer exchanged into a storage buffer (10mM KPO_4_^-^ pH 6.8, 10 mM DTT). Purified PCK2 was concentrated (ε_280_=16.4mL/mg), flash-frozen in 30 μl drops by direct immersion into liquid nitrogen and subsequently stored at −80°C.

### Sample preparation for Biotin Thiol Assay

Sample preparation and analysis were based on (Gao et al., 2015). In brief, proteins were extracted from cells with RIPA buffer (150 mM NaCl, 1 mM EDTA, 0.5% Triton X-100, 0.5% deoxycholic acid, and 100 mM Tris pH 7.5) containing protease and phosphatase inhibitors. 2 mg of protein was incubated with 100 μM NM-biotin (Pierce) for 30min, then mixed with Streptavidin-agarose resin (Thermo Scientific) and kept rotating overnight at 4°C. The beads were washed and eluted with DTT or TCEP (10 mM). Eluted proteins were concentrated to a final volume of 25–40 μl with using of Amicon Ultracel 10K (Millipore), and used for gel electrophoresis followed by western blot analysis.

### Sample preparation for TMT-BTA assay

Proteins (2 mg) were extracted and biotinylated as described above. Biotinylated proteins were precipitated with ice cold acetone, resuspended in denaturation buffer (8 M urea, 1 mM MgSO_4_ and 30 mM Tris–HCl, pH 7.5) diluted with 10 volumes of buffer containing 1 mM CaCl_2_, 100 mM NaCl and 30 mM HEPES-NaOH pH 8.0, then incubated with MS-grade trypsin (Thermo, 90058) with occasional mixing for 18h at 37°C. The ratio of the enzyme to substrate was 1:40 (w/w). After digestion, trypsin was inactivated by incubation at 95°C for 10min, then reactions were mixed with streptavidin-agarose beads (0.5-2 ml) and incubated at 4°C for 18h following extensive washes in the presence of 0.1% SDS. Peptides were eluted with the buffer containing 10 mM TEAB with 10 mM TCEP. TCEP was removed with using a C-18 column (Thermo). Peptides were eluted from the desalting column with 80% methanol, dried under vacuum, and suspended in buffer (50 mM HEPES-NaOH pH 8.0). Free −SH groups were alkylated by iodoTMT reagents at final volume of 150 μl. After 1h of TMT labeling, addition of DTT (20 mM) terminated the alkylation reaction. After desalting through C-18 columns, the alkylated peptides were mixed and combined for LC-MS analysis.

### TMT-BTA peptide identification and quantification by LC-MS analysis

This iodoTMT 6plex labeled peptide sample was reconstituted into 30 μl 1% acetic acid and was ready for MS analysis. The LC-MS system was a Thermo Ultimate 3000 UHPLC interfaced with a ThermoFisher Scientific Fusion Lumos tribrid mass spectrometer system. The HPLC column was a Dionex 15 cm x 75 μm id Acclaim Pepmap C18, 2μm, 100 Å reversed-phase capillary chromatography column. Five μl volumes of the extract were injected, and the peptides eluted from the column by an acetonitrile/0.1% formic acid gradient at a flow rate of 0.3 μl/min were introduced into the source of the mass spectrometer on-line. The nanospray ion source was operated at 1.9 kV.

The digest was analyzed using both TMT-MS2 method and TMT-MS3 method. The TMT-MS2 is a data dependent acquisition method using HCD fragmentation for MS/MS scans of the precursor ions selected from MS1 full scan. The MS2 spectra were used for peptide identifications and the low mass region (100-140) of the same spectra was used simultaneously for quantifications of these peptides.

The TMT-MS3 method uses a technique called synchronized precursor selection (SPS) that can only be achieved on Thermo Fusion series MS. The method is also a data dependent acquisition using CID fragmentation for the MS2 scan of precursors selected from MS1. Then consecutive MS3 HCD scans at 100-500 m/z were performed on a combination of several of the most abundant ions (set at 10 in this study) selected from the MS2 scans. The CID MS2 spectra were used for peptide identifications and the HCD MS3 spectra were used for peptide quantifications.

### Label-free MS quantification of full proteome

The gel contains two samples derived from IHH and EndoC-BH3 cells. Each gel lane was cut into eight areas, these areas were digested with trypsin, and the digests were analyzed by LC-MS as described above. The data was queried against the human UniProtKB database. Protein quantitative value expressed as LFQ intensity was generated by precursor intensity based label-free quantification algorithm used by MaxQuant. LFQ intensities are the output of the Max-LFQ algorithm (Cox et al., 2014). These are based on the raw intensities that are normalized on multiple levels to ensure that profiles of LFQ intensities across samples accurately reflect the relative amounts of the proteins. Two different filters were used to identify proteins that have a different abundance in these samples, including a two-fold difference in LFQ values, and a minimum of at least 5 peptides.

### Immunoprecipitation of of human FLAG-PCK2

MIN6 cells expressing FLAG-tagged human PCK2 were washed with ice-cold PBS and harvested. The cells were lysed in RIPA lysis buffer (150 mM NaCl, 1 mM EDTA, 0.5% Triton X-100, 0.5% deoxycholic acid, 5 mM NEM and 100 mM Tris–HCl pH 7.5) containing protease and phosphatase inhibitors. 4 mg of protein extracts were incubated with Anti FLAG M2 affinity gels (50 μl, Sigma) with rotation overnight at 4°C. The gels were washed once with RIPA buffer and twice with HBS buffer (100 mM NaCl, 50 mM HEPES-NaCl). The FLAG-tagged PCK2 proteins were eluted with 100 μl of 3X FLAG peptides at 0.15 μg/μl. Immunoprecipitated proteins were resolved by a nonreducing SDS-PAGE.

### LC-MS identification of PCK2 S-sulfhydration at the catalytic cysteine 306 residue

Human FLAG-PCK2 was expressed, and purified as described above in MIN6 cells treated with Tg for 18h. The protein band in the gel was digested in-gel by adding 10 μl 5 ng/μl trypsin in 50 mM ammonium bicarbonate and incubating overnight at room temperature to achieve complete digestion. The peptides that were formed were extracted from the polyacrylamide in two aliquots of 30 μl 50% acetonitrile with 5% formic acid. These extracts were combined and evaporated to <10 μl in Speedvac and analyzed directly by LC-MS. HPLC was performed as described previously. The microelectrospray ion source was operated at 2.5 kV. The digest was analyzed using the data dependent multitask capability of the instrument acquiring full scan mass spectra to determine peptide molecular weights and product ion spectra to determine amino acid sequence in successive instrument scans. The data were analyzed by using all CID spectra collected in the experiment to query the human UniProtKB databases with the search programs Mascot and more specifically against the sequence of PCK2 with the Sequest program.

### LC-MS identification of human recombinant insulin S-sulfhydration

22.2 μM human recombinant insulin was incubated with 25 μg CTH in the presence of 4 mM L-cysteine and 1mM PLP in buffer (100 mM HEPES-NaOH, pH7.4) for 1h. The reaction mixture was dried and submitted for a LC-MS analysis.

### Generation of CIRSPR-mediated knockout CTH INS1 cell lines

CTH null INS1 cells were generated using the protocol described in (Sanjana et al., 2014). In brief, sgRNAs targeting rat CTH were designed, amplified and cloned into transient lentiCRISPR v2 (Addgene plasmid # 52961) as a gift from Feng Zhang. Mammalian lentiviral particles harboring sgRNA-CTH plasmids were generated as described in (Sanjana et al., 2014). After 3 days of infection of INS1 cells, puromycin was added to enrich Cas9-expressing cells. Following 10 days of the puromycin selection, individual clone were isolated by flow cytometry and analyzed for decreased expression of CTH when overexpressed ATF4.

sgRNA sequences for rat CTH gene:

sgCTH1: 5’-GGAGGGCGTCCTTCTGCATG-3’

sgCTH2: 5’-GAGGCGTCCTTCTGCATGCT-3’

### Mismatch detection assay

Genomic DNA was extracted from individual clone INS1 cells using a DNA extraction kit (Zeymo). Target regions were PCR-amplified with *pfu* DNA polymerase (phusion, NEB, M0530S). PCR products were denatured at 95°C for 10min and re-annealed at −2°C/second temperature ramp to 85°C, followed by a −1 °C/second ramp to 25°C. The heterocomplexed PCR products (5 μl) were incubated with 5 U T7E1 enzyme (New England Bio Labs) at 37°C for 20min. Products from mismatch assays were measured by electrophoresis on 2% agarose gel. Two different PCR primers (set 1 and set 2) were used to amplify the regions flanking the CRISPR targeting sites.

Primer 1 (F): 5’-GCTTCCTCATCCTGCAGACA-3’

Primer 1 (R): 5’-CCTGCTTACAGCCTGATCTTCT-3’

Primer 2 (F): 5’-CTCACCGGCTTCCTCATCCT-3’

Primer 2 (R): 5-GCCTGATCTTCTGGAGGCATTT-3’

### Measurement of recombinant PCK2 activity in vitro

A continuous enzymatic assay was used to measure PCK activity, in which OAA produced by PCK is immediately reduced to malate in a coupled assay with malate dehydrogenase as described by (Johnson and Holyoak, 2012). The reactions were performed in triplicate in 96-well plates with a final volume of 200 μl containing 50 mM HEPES (pH 7.5), 2 mM MgCl_2_, 500 μM GTP, 1 mM ADP, 2 mM MnCl_2_, 150 μM NADH, and 8 units/ml of malic dehydrogenase (Sigma). The reaction was initiated with 0.5 mM PEP (Sigma), and the decrease in NADH signal was assayed for 3 min at 3 second intervals by absorbance at 340 nm (M3 plate reader, Molecular Devices).

### Identification of protein S-glutathionylation by the Biotin Switch Technique (BST)

Similar to the BST used for the detection of nitrosylated proteins, detection of oxidized proteins from protein extracts was performed using the BST following the procedure described in (Gao et al., 2015) with minor modifications. Briefly, protein extraction was performed as described above. After alkylation with NEM, samples were precipitated with three volumes of cold acetone at −20°C for 30min to remove excess alkylating reagents, centrifuged (15,000 g, 10min, 4°C), and resuspended at a concentration of 1 mg/ml in solubilization buffer (30 mM Tris-HCl pH 7.9, 1 mM EDTA and 100 mM NaCl) supplemented with 0.5% SDS. Reduction of oxidized proteins was achieved by adding 10 mM TCEP, then followed by cold acetone precipitation to remove excess TCEP. TCEP-treated protein pellets were dissolved in the solubilization buffer, then mixed with 0.25 mM HPDP-Biotin to label free thiols. After 60 min incubation at room temperature, the excess of HPDP-biotin was removed by acetone precipitation, and precipitated proteins were then resuspended at 1 mg/ml in solubilization buffer. The biotinylated sample was incubated with streptavidin beads (Thermo-20357) rotating at 4°C for overnight. The beads were then washed 6 times with wash buffer (600 mM NaCl, 1 mM EDTA and 50 mM HEPES-NaOH, pH7.5) and eluted with TCEP. Protein eluate was separated by non-reducing SDS-PAGE following Western blot with the indicated antibodies.

### Metabolic labeling of cells with [^5^S] Met

Sample preparation and analysis were based on (Guan BJ et al, JBC 2014). Briefly, INS1 cells were exposed to diamide at the indiciated concentrations for 1h, then washed with warm PBS. The untreated and treated INS1 cells incubated for 10 min in Met/Cys-free RPMI supplemented with 10% heat-inactive FBS in the presence of the treatment reagents. [^35^S]Met was added (30 μCi/ml) for an additional 30 min. INS1 cells were rinsed twice with cold PBS, and total proteins were precipitated three times with 5% TCA and 1 mM Met for 10 min on ice. Precipitates were dissolved in 200 μl of 1 N NaOH and 0.5% sodium deoxycholate for 1 h. Radioactivity was determined by liquid scintillation counting. Total cellular proteins were quantified by the DC^TM^ Protein Assay (Bio-Rad) following the manufacturer’s instruction. Incorporation of [^35^S] Met/Cys into total cellular proteins was calculated and normalized.

### Metabolite analysis by targeted LC-MS

Determination of glucose flux:

Treatment and assay of media [U-^13^ C_6_] glucose enrichment

INS1 cells were seeded and cultured in the cell growth medium (RPMI, 1640 Thermo) containing 11.1 mM glucose, 10% heat-inactive FBS, 1 mM sodium pyruvate, 2 mM glutamine, 10 mM HEPES, 100 U/ml penicillin and 100 μg/ml streptomycin. Media were refreshed after 48h. INS1 cells were washed with warm PBS for one time, then incubated with glucose-free KRB buffer in the presence or absence of 0.1 mM diamide for 90 min. The untreated and treated cells were then exposed to the KRB buffer (a mixture of 8.4 mM glucose plus 8.4 mM of [U-^13^C_6_] glucose, and the indicated concentrations of H_2_S donor). After 90 min incubation, the cells were scraped and pellets were stored at −80°C until extraction of metabolites.

Metabolite extraction: Metabolites were extracted from cell pellets as described (Gao et al., 2015).

GC-MS conditions: Analyses were carried out on an Agilent 5973 mass spectrometer equipped with 6890 Gas Chromatograph. The details of GC-MS were described (Gao et al., 2015).

Determination of PEP flux:

Treatment and assay of media [M+1-^13^C] PEP enrichment

MIN6 cells were plated onto 10-cm plates and cultured in the cell growth medium. After 48h, metabolic labeling was performed. Cells were washed with warm PBS and incubated for 3.5h in the DMEM medium containing 10% heat inactive FBS, 2 mM glutamine, 2 mM sodium pyruvate [3-^13^C_1_], 100 U/ml penicillin and 100 μg/ml streptomycin and 25 mM glucose. After the incubation, MIN6 cells were scraped and pellets were stored at −80°C until extraction of metabolites.

Metabolite extraction: Metabolites were extracted from cell pellets as described (Stark et al., 2009).

LC-MS conditions: Analyses were carried out on a Shimadzu 8040 mass spectrometer. The details of LC-MS were described (Stark et al., 2009).

## References

Anastasiou, D., Poulogiannis, G., Asara, J.M., Boxer, M.B., Jiang, J.K., Shen, M., Bellinger, G., Sasaki, A.T., Locasale, J.W., Auld, D.S., et al. (2011). Inhibition of pyruvate kinase M2 by reactive oxygen species contributes to cellular antioxidant responses. Science 334, 1278–1283.

Cao, S.S., and Kaufman, R.J. (2014). Endoplasmic reticulum stress and oxidative stress in cell fate decision and human disease. Antioxidants& redox signaling 21, 396–413.

Cox, J., Hein, M.Y., Luber, C.A., Paron, I., Nagaraj, N., and Mann, M. (2014). Accurate proteome-wide label-free quantification by delayed normalization and maximal peptide ratio extraction, termed MaxLFQ. Molecular& cellular proteomics : MCP 13, 2513–2526.

Das, A., Huang, G.X., Bonkowski, M.S., Longchamp, A., Li, C., Schultz, M.B., Kim, L.J., Osborne, B., Joshi, S., Lu, Y, et al. (2018). Impairment of an Endothelial NAD(+)-H2S Signaling Network Is a Reversible Cause of Vascular Aging. Cell 173, 74–89 e20.

Filipovic, M.R., Zivanovic, J., Alvarez, B., and Banerjee, R. (2018). Chemical Biology of H2S Signaling through Persulfidation. Chemical reviews 118, 1253–1337.

Fratelli, M., Demol, H., Puype, M., Casagrande, S., Eberini, I., Salmona, M., Bonetto, V., Mengozzi, M., Duffieux, F., Miclet, E, et al. (2002). Identification by redox proteomics of glutathionylated proteins in oxidatively stressed human T lymphocytes. Proceedings of the National Academy of Sciences of the United States of America 99, 3505–3510.

Gao, X.H., Krokowski, D., Guan, B.J., Bederman, I., Majumder, M., Parisien, M., Diatchenko, L., Kabil, O., Willard, B., Banerjee, R., et al. (2015). Quantitative H2S-mediated protein sulfhydration reveals metabolic reprogramming during the integrated stress response. eLife 4, e10067.

Guan, B.J., van Hoef, V., Jobava, R., Elroy-Stein, O., Valasek, L.S., Cargnello, M., Gao, X.H., Krokowski, D., Merrick, W.C., Kimball, S.R., et al. (2017). A Unique ISR Program Determines Cellular Responses to Chronic Stress. Molecular cell 68, 885–900 e886.

Gerber, P.A., and Rutter, G.A. (2017). The Role of Oxidative Stress and Hypoxia in Pancreatic Beta-Cell Dysfunction in Diabetes Mellitus. Antioxidants& redox signaling 26, 501–518.

Haataja, L., Snapp, E., Wright, J., Liu, M., Hardy, A.B., Wheeler, M.B., Markwardt, M.L., Rizzo, M., and Arvan, P. (2013). Proinsulin intermolecular interactions during secretory trafficking in pancreatic beta cells. The Journal of biological chemistry 288, 1896–1906.

Han, J., Back, S.H., Hur, J., Lin, Y.H., Gildersleeve, R., Shan, J., Yuan, C.L., Krokowski, D., Wang, S., Hatzoglou, M., et al. (2013). ER-stress-induced transcriptional regulation increases protein synthesis leading to cell death. Nature cell biology 15, 481–490.

Hine, C., Harputlugil, E., Zhang, Y., Ruckenstuhl, C., Lee, B.C., Brace, L., Longchamp, A., Trevino-Villarreal, J.H., Mejia, P., Ozaki, C.K., et al. (2015). Endogenous hydrogen sulfide production is essential for dietary restriction benefits. Cell 160, 132–144.

Hiranruengchok, R., and Harris, C. (1995). Diamide-induced alterations of intracellular thiol status and the regulation of glucose metabolism in the developing rat conceptus in vitro. Teratology 52, 205–214.

Johnson, T.A., and Holyoak, T. (2012). The Omega-loop lid domain of phosphoenolpyruvate carboxykinase is essential for catalytic function. Biochemistry 51, 9547–9559.

Kabil, H., Kabil, O., Banerjee, R., Harshman, L.G., and Pletcher, S.D. (2011). Increased transsulfuration mediates longevity and dietary restriction in Drosophila. Proceedings of the National Academy of Sciences of the United States of America 108, 16831–16836.

Kebede, M., Favaloro, J., Gunton, J.E., Laybutt, D.R., Shaw, M., Wong, N., Fam, B.C., Aston-Mourney, K., Rantzau, C., Zulli, A., et al. (2008). Fructose-1,6-bisphosphatase overexpression in pancreatic beta-cells results in reduced insulin secretion: a new mechanism for fat-induced impairment of beta-cell function. Diabetes 57, 1887–1895.

Li, L., Rose, P., and Moore, P.K. (2011). Hydrogen sulfide and cell signaling. Annual review of pharmacology and toxicology 51, 169–187.

Liu, M., Hodish, I., Haataja, L., Lara-Lemus, R., Rajpal, G., Wright, J., and Arvan, P. (2010). Proinsulin misfolding and diabetes: mutant INS gene-induced diabetes of youth. Trends Endocrinol Metab 21, 652–659.

Longchamp, A., Mirabella, T., Arduini, A., MacArthur, M.R., Das, A., Trevino-Villarreal, J.H., Hine, C., Ben-Sahra, I., Knudsen, N.H., Brace, L.E., et al. (2018). Amino Acid Restriction Triggers Angiogenesis via GCN2/ATF4 Regulation of VEGF and H2S Production. Cell 173, 117–129 e114.

Luo, W., and Semenza, G.L. (2012). Emerging roles of PKM2 in cell metabolism and cancer progression. Trends in endocrinology and metabolism: TEM 23, 560–566.

Mailloux, R.J., Jin, X., and Willmore, W.G. (2014). Redox regulation of mitochondrial function with emphasis on cysteine oxidation reactions. Redox biology 2, 123–139.

Martinez-Ruiz, A., and Lamas, S. (2007). Signalling by NO-induced protein S-nitrosylation and S-glutathionylation: convergences and divergences. Cardiovascular research 75, 220–228.

Mann, M., and Jensen, O.N. (2003). Proteomic analysis of post-translational modifications. Nature biotechnology 21, 255–261.

Mieyal, J.J., Gallogly, M.M., Qanungo, S., Sabens, E.A., and Shelton, M.D. (2008). Molecular mechanisms and clinical implications of reversible protein S-glutathionylation. Antioxidants& redox signaling 10, 1941–1988.

Mironov, A., Seregina, T., Nagornykh, M., Luhachack, L.G., Korolkova, N., Lopes, L.E., Kotova, V., Zavilgelsky, G., Shakulov, R., Shatalin, K., et al. (2017). Mechanism of H2S-mediated protection against oxidative stress in Escherichia coli. Proceedings of the National Academy of Sciences of the United States of America 114, 6022–6027.

Mishanina, T.V., Libiad, M., and Banerjee, R. (2015). Biogenesis of reactive sulfur species for signaling by hydrogen sulfide oxidation pathways. Nat Chem Biol 11, 457–464.

Miller, D.L., and Roth, M.B. (2007). Hydrogen sulfide increases thermotolerance and lifespan in Caenorhabditis elegans. Proceedings of the National Academy of Sciences of the United States of America 104, 20618–20622.

Mustafa, A.K., Gadalla, M.M., Sen, N., Kim, S., Mu, W., Gazi, S.K., Barrow, R.K., Yang, G., Wang, R., and Snyder, S.H. (2009). H2S signals through protein S-sulfhydration. Science signaling 2, ra72.

Paul, B.D., and Snyder, S.H. (2012). H(2)S signalling through protein sulfhydration and beyond. Nature reviews Molecular cell biology 13, 499–507.

Paul, B.D., Sbodio, J.I., Xu, R., Vandiver, M.S., Cha, J.Y., Snowman, A.M., and Snyder, S.H. (2014). Cystathionine gamma-lyase deficiency mediates neurodegeneration in Huntington’s disease. Nature 509,96–100.

Paul, B.D., and Snyder, S.H. (2015). H2S: A Novel Gasotransmitter that Signals by Sulfhydration. Trends in biochemical sciences 40, 687–700.

Reddie, K.G., and Carroll, K.S. (2008). Expanding the functional diversity of proteins through cysteine oxidation. Current opinion in chemical biology 12, 746–754.

Sanjana, N.E., Shalem, O., and Zhang, F. (2014). Improved vectors and genome-wide libraries for CRISPR screening. Nature methods 11, 783–784.

Sbodio, J.I., Snyder, S.H., and Paul, B.D. (2018). Regulators of the transsulfuration pathway. British journal of pharmacology.

Sen, N., Paul, B.D., Gadalla, M.M., Mustafa, A.K., Sen, T., Xu, R., Kim, S., and Snyder, S.H. (2012). Hydrogen sulfide-linked sulfhydration of NF-kappaB mediates its antiapoptotic actions. Molecular cell 45,13–24.

Stark, R., Pasquel, F., Turcu, A., Pongratz, R.L., Roden, M., Cline, G.W., Shulman, G.I., and Kibbey, R.G. (2009). Phosphoenolpyruvate cycling via mitochondrial phosphoenolpyruvate carboxykinase links anaplerosis and mitochondrial GTP with insulin secretion. The Journal of biological chemistry 284, 26578–26590.

Szabo, C. (2012). Roles of hydrogen sulfide in the pathogenesis of diabetes mellitus and its complications. Antioxidants& redox signaling 17, 68–80.

Szabo, C., Coletta, C., Chao, C., Modis, K., Szczesny, B., Papapetropoulos, A., and Hellmich, M.R. (2013). Tumor-derived hydrogen sulfide, produced by cystathionine-beta-synthase, stimulates bioenergetics, cell proliferation, and angiogenesis in colon cancer. Proceedings of the National Academy of Sciences of the United States of America 110, 12474–12479.

Thompson, A., Schafer, J., Kuhn, K., Kienle, S., Schwarz, J., Schmidt, G., Neumann, T., Johnstone, R., Mohammed, A.K., and Hamon, C. (2003). Tandem mass tags: a novel quantification strategy for comparative analysis of complex protein mixtures by MS/MS. Analytical chemistry 75, 1895–1904.

Ulatowski, L., Dreussi, C., Noy, N., Barnholtz-Sloan, J., Klein, E., and Manor, D. (2012). Expression of the alpha-tocopherol transfer protein gene is regulated by oxidative stress and common single-nucleotide polymorphisms. Free radical biology& medicine 53, 2318–2326.

Vander Heiden, M.G., Cantley, L.C., and Thompson, C.B. (2009). Understanding the Warburg effect: the metabolic requirements of cell proliferation. Science 324, 1029–1033.

Yang, W., Yang, G., Jia, X., Wu, L., and Wang, R. (2005). Activation of KATP channels by H2S in rat insulin-secreting cells and the underlying mechanisms. The Journal of physiology 569, 519–531.

Yang, G., Yang, W., Wu, L., and Wang, R. (2007). H2S, endoplasmic reticulum stress, and apoptosis of insulin-secreting beta cells. The Journal of biological chemistry 282, 16567–16576.

Yang, G., Wu, L., Jiang, B., Yang, W., Qi, J., Cao, K., Meng, Q., Mustafa, A.K., Mu, W., Zhang, S., et al.(2008). H2S as a physiologic vasorelaxant: hypertension in mice with deletion of cystathionine gamma-lyase. Science 322, 587–590.

van der Linde, K., Gutsche, N., Leffers, H.M., Lindermayr, C., Muller, B., Holtgrefe, S., and Scheibe, R. (2011). Regulation of plant cytosolic aldolase functions by redox-modifications. Plant physiology and biochemistry : PPB 49, 946–957.

Wan, X., Zinselmeyer, B.H., Zakharov, P.N., Vomund, A.N., Taniguchi, R., Santambrogio, L., Anderson, M.S., Lichti, C.F., and Unanue, E.R. (2018). Pancreatic islets communicate with lymphoid tissues via exocytosis of insulin peptides. Nature 560, 107–111.

Zaffagnini, M., Bedhomme, M., Groni, H., Marchand, C.H., Puppo, C., Gontero, B., Cassier-Chauvat, C., Decottignies, P., and Lemaire, S.D. (2012). Glutathionylation in the photosynthetic model organism Chlamydomonas reinhardtii: a proteomic survey. Molecular& cellular proteomics : MCP 11, M111 014142.

